# Robustness and reproducibility of simple and complex synthetic logic circuit designs using a DBTL loop

**DOI:** 10.1101/2022.06.10.495560

**Authors:** Breschine Cummins, Justin Vrana, Robert C. Moseley, Hamed Eramian, Anastasia Deckard, Pedro Fontanarrosa, Daniel Bryce, Mark Weston, George Zheng, Joshua Nowak, Francis C. Motta, Mohammed Eslami, Kara Layne Johnson, Robert P. Goldman, Chris J. Myers, Tessa Johnson, Matthew W. Vaughn, Niall Gaffney, Joshua Urrutia, Shweta Gopaulakrishnan, Vanessa Biggers, Trissha R. Higa, Lorraine A. Mosqueda, Marcio Gameiro, Tomáš Gedeon, Konstantin Mischaikow, Jacob Beal, Bryan Bartley, Tom Mitchell, Tramy T. Nguyen, Nicholas Roehner, Steven B. Haase

**Affiliations:** Montana State University, Bozeman, MT, U.S.A.; UW BIOFAB, Seattle, WA, U.S.A.; Duke University, Durham, NC, U.S.A.; Netrias LLC, Annapolis, MD, U.S.A.; Geometric Data Analytics, Inc., Durham, NC, U.S.A.; University of Utah, Salt Lake City, UT, U.S.A.; Sift, LLC, Minneapolis, MN, U.S.A.; Strateos, Inc., Menlo Park, CA, U.S.A.; Florida Atlantic University, Boca Raton, FL, U.S.A.; University of Colorado Boulder, Boulder, CO, U.S.A.; Texas Advanced Computing Center, Austin, TX, U.S.A.; Rutgers University, New Brunswick, NJ, U.S.A.; Raytheon BBN, Cambridge, MA, U.S.A.

**Keywords:** Synthetic logic circuits, Design-Build-Test-Learn, flow cytometry, CRISPR, yeast, genetic circuits, machine learning, automated experiments

## Abstract

Computational tools addressing various components of design-build-test-learn loops (DBTL) for the construction of synthetic genetic networks exist, but do not generally cover the entire DBTL loop. This manuscript introduces an end-to-end sequence of tools that together form a DBTL loop called DART (Design Assemble Round Trip). DART provides rational selection and refinement of genetic parts to construct and test a circuit. Computational support for experimental process, metadata management, standardized data collection, and reproducible data analysis is provided via the previously published Round Trip (RT) test-learn loop. The primary focus of this work is on the Design Assemble (DA) part of the tool chain, which improves on previous techniques by screening up to thousands of network topologies for robust performance using a novel robustness score derived from dynamical behavior based on circuit topology only. In addition, novel experimental support software is introduced for the assembly of genetic circuits. A complete design-through-analysis sequence is presented using several OR and NOR circuit designs, with and without structural redundancy, that are implemented in budding yeast. The execution of DART tested the predictions of the design tools, specifically with regard to robust and reproducible performance under different experimental conditions. The data analysis depended on a novel application of machine learning techniques to segment bimodal flow cytometry distributions. Evidence is presented that, in some cases, a more complex build may impart more robustness and reproducibility across experimental conditions.

## 1. Introduction

The construction of synthetic biology genetic circuits is a growing field that holds great promise for producing purpose-built cellular machines that perform important tasks, such as monitoring environmental conditions and producing materials or therapeutics (1; 2; 3). The reproducibility of these constructs is of critical importance. The most common approaches to achieving reproducibility in results seek to standardize protocols and analyses and tightly control experimental conditions. However, in the field of synthetic biology, reproducible function of synthetic genetic circuits may be closely tied to the robustness of the construct design (4). By incorporating measures of robustness into the design principles of synthetic biology, one may be able to generate constructs that have functions that are robust to changes in genetic components or in experimental conditions, causing increased reproducibility across laboratories.

There are many stages in the design of synthetic constructs (5), including: (i) the choice of circuit structure (also called topology in this work), (ii) the DNA sequence design of a genetic parts library, (iii) the choice of genetic parts (taken from the parts library) used within the modular circuit structure, (iv) the choice of insertion points or landing pads in the genome or plasmid, and (v) the design of the experimental protocol used to assess the resulting design. All of these stages are facilitated by the rational application of computational tools to reduce experimental time and effort and to improve success rate.

Many computational tools are available for different parts of the design process. Examples include methods for DNA sequence design of genetic parts (6; 7), for parts choice and construction into a linear DNA sequence (8; 9; 10), for landing pad choice (11), and for circuit structure or topology (12). Experimental protocols are generally custom designed for each application, but tools exist to facilitate the reproducibility of protocols (13; 14). Some tools integrate several stages of the design process together; for example, the software tool Cello (15; 16) takes a logic function written in the Verilog language and identifies a single circuit structure using design principles from electronic circuit design that employ NOT and NOR gates. Cello then optimizes the modular construction of the logic circuit from a parts library to create a linear DNA sequence with the desired circuit. Another method (17) exhaustively explores fan-out free circuit structures and jointly optimizes structure and parts assignment for a specific logic function.

Design methods are validated by the construction of the predicted optimal circuit design(s), which are then analyzed for performance. The conjunction of the design stage with the build, experimentation, and analysis of a synthetically built genetic circuit forms a design-build-test-learn (DBTL) loop (18), in which the results of the analysis can be used to refine the experimental protocol or to tweak the design. The need for end-to-end tooling of DBTL loops, particularly when high-throughput data generation is available, is recognized (19).

In this work, a design-build-test-learn loop called DART (Design Assemble Round Trip) is presented for the rational design of synthetic biology genetic logic circuits. In principle the technology is generalizable to dynamically-complex circuit functions beyond logic (20) that are of interest to the synthetic biology community (21; 22; 23). DART is comprised of tools for (i) the prediction of robust circuit topologies, (ii) prediction of the most effective choice of parts to construct the topology, (iii) sequence construction for selected designs, (iv) step-by-step instructions for build assembly, and (v) reproducible experimental submission, data and metadata consolidation, data standardization, and automated data analysis using a previously published test-learn loop called the Round Trip (RT).

The Design Assemble (DA) part of the tool chain is the primary focus in this work. DA, like Cello, starts with a logic function and a library of genetic parts characterized by dosage-response curves and ultimately produces a linear DNA sequence. The design stage differs from Cello by first screening alternative network topologies—possibly thousands depending on the allowed maximum size of the circuit—and scoring them for dynamically robust performance. Using a similar approach, it was previously shown in (17) that varying circuit topology can result in improved performance. Our approach differs from (17) by the choice of a robustness score based solely on circuit topology that reduces the number of parts optimizations required. Once the most promising circuit topologies are chosen by the design stage of DA, parts are assigned using a machine learning technique and assembled into a linear sequence. Automated lab software then increases the reproducibility of the build stage.

Ideally, a genetic construct will function robustly under a variety of conditions, since in practice it can be difficult to reproduce experiments across labs (24). Robust genetic constructs make it easier to achieve reproducibility, since they make design performance less susceptible to differences between lab conditions. The primary purpose of DART is to produce synthetic logic circuit designs embedded in cells that perform adequately under the widest possible range of conditions and provide reproducible and consistent results.

### 1.1. Software tool chain

DART provides a systematic and standardized approach to building genetic constructs with desired functional properties. A computational framework was developed that supports the design and construction of synthetic logic circuits from a library of dosage-response characterized parts, DA (Design Assemble), and was attached to an existing tool chain RT (Round Trip, (25; 26)) that standardizes data collection, preprocessing, and analysis. A diverse set of tools was identified, collected, and unified to meet the needs of cellular circuit construction from design to analysis. See Fig. 1 for a schematic of the connections between the tools.

**Fig. 1.**
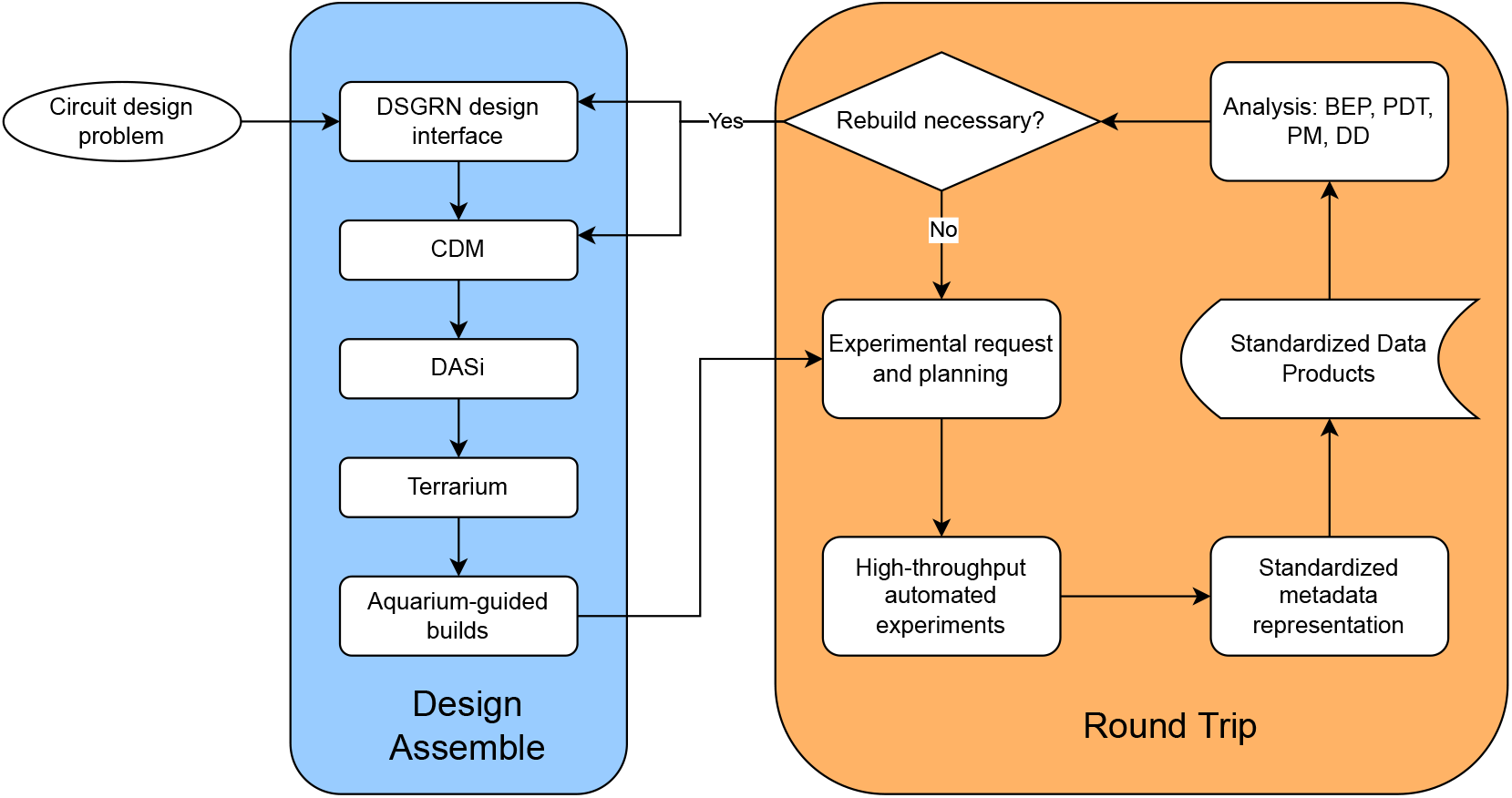
Flow chart showing the details of DART, particularly the integration between Design Assemble (introduced here) and Round Trip (25).

The RT tool chain supports experimental metadata development and maintenance by automating tedious metadata design and encoding and by reacting and repairing as deviations arise. The RT connects experimental data and subsequent analyses with the deeply-represented experimental DA constructs via user-friendly construct names. For example, RT users develop experimental requests referencing the common names for the constructs developed by DA, which are automatically resolved against SBOL (27; 28) representations of the DA constructs. The RT carries these resolved references through the experimental process to link constructs to experimental results. The resulting data and metadata represents a rich, AI-ready data set that the RT can automatically analyze and present or package for alternative analysis tools.

The design and build aspects of DART have not been previously used in combination. The design/prediction tools are Dynamic Signatures Generated by Regulatory Networks (DSGRN, (20; 29; 30)) and Combinatorial Design Model (CDM (31; 32)). The build tools are DNA Assembly (DASi, (33; 34; 35)), the computer-aided process planning tool Terrarium (36; 35), and the lab software Aquarium (14; 37).

DSGRN is a flexible and highly scalable tool for analyzing and predicting all possible long term dynamics of a regulatory network. It requires only a network topology with annotations indicating whether regulatory interactions are activating or repressing. The computed dynamics exhaustively describe the possible long-term behaviors that the network is capable of exhibiting. Although used here to assess the equilibrium values of logic circuits, DSGRN is not limited to modeling and analysis of logic functions; its capabilities greatly exceed that specialized problem. Its integration into DA involved the implementation of a user-friendly DSGRN Design Interface (38) dedicated to the express purpose of designing synthetic logic circuits. The DSGRN Design Interface incorporates qualitative build constraints in a plain language input file. Interpretation of the output circuit topology scoring is readily accessible via figures and human-readable descriptions of constraints on the interaction of genetic parts, and there is also the option of output in machine readable SBOL2 format (27).

CDM is a neural-network based model that makes in-silico predictions of experiments by using context and data from a subset of conditions and predicts the outcome in all combinations of conditions. The application of CDM in this work optimized a combination of genetic parts for a given circuit using training data from single parts.

Terrarium bridges the gap between synthetic biology design specification and the build process through computer-aided process planning. A biological design is encoded as a biological manufacturing file (BMF). Terrarium converts BMFs into executable networks of protocols, called workflows. These workflows are uploaded to the laboratory software Aquarium, which manages laboratory inventory, automates protocol execution, and generates human-readable instructions to execute the workflows. Terrarium has an internal digital model of the laboratory that is periodically updated by metadata collected in Aquarium, specifically protocol execution time, inventory usage, experimental errors, success rates, materials, and labor costs. Using this model, Terrarium can predict lead times and costs that inform future workflow process planning for increased accuracy, economy, and efficiency. DASi is a subordinate algorithm of Terrarium released as stand-alone software that provides assembly instructions for a DNA sequence when that sequence does not already exist in lab inventory. It produces a BMF that is ingested by Terrarium to create a workflow for Aquarium.

### 1.2. Biological Scenario

The main hypothesis is that the benefits of genetic network redundancy can outweigh the costs of circuit build complexity through increased reproducibility and robustness across experimental conditions. A network with built-in redundancy and parts optimization has the capacity to exhibit better performance (17), but increased complexity leads to more difficult builds.

Circuit robustness was examined by comparing the performance of simple and complex network topologies of OR and NOR synthetic circuits in the yeast *Saccharomyces cerevisiae*, while also testing the predictions for two different sets of parts for each topology using CDM. The performance of a circuit is evaluated as a circuit’s ability to express fluorescence (ON) or not (OFF) as intended by the circuit’s logic given the presence or absence of chemical inputs. Logic circuits designed to exhibit OR and NOR logic were chosen based on preliminary data analysis in which previously built OR and NOR circuits performed poorly ((4; 39) and Fig. 6 in (25)). See Fig. 2 for schematics of the designs discussed in this work, where each quadrant shows one topology with two CDM designs. The top row shows the simplest topologies that are capable of producing the desired circuit behavior. The alternative topologies discovered using the DSGRN Design Interface are called DSGRN topologies and shown in the second row.

**Fig. 2.**
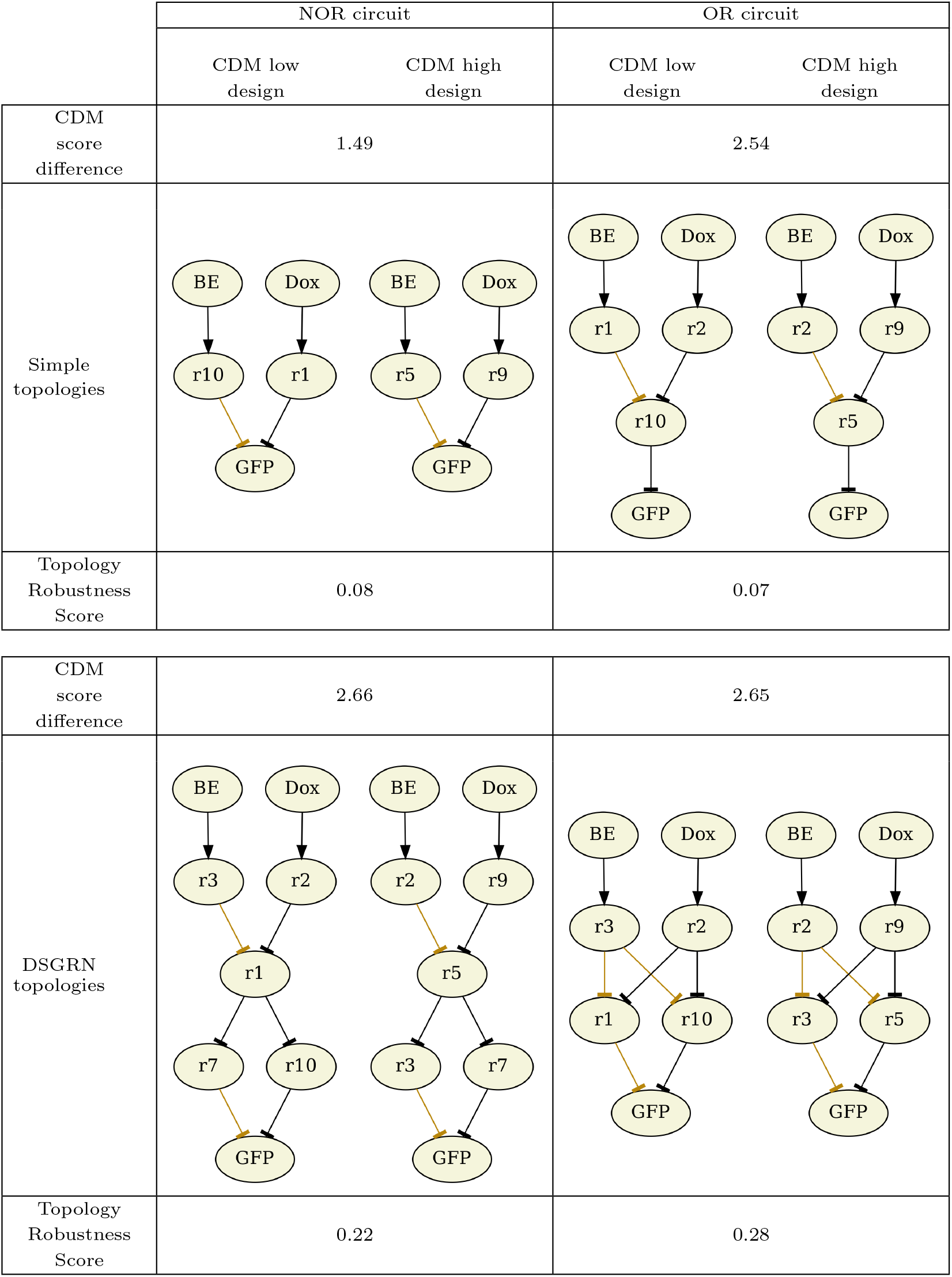
The designs for the built circuits, with one topology and two CDM designs in each quadrant. Upper left: The simple NOR topology published in (40), re-engineered with two different collections of gRNA inducible parts. The one on the left is predicted by CDM to perform worse than the one on the right. Lower left: The DSGRN NOR circuit predicted by DSGRN to perform more robustly than the simple NOR circuit, with the two CDM predicted gRNA arrangements. Upper right: The simple OR topology published in (40), with CDM-selected parts assignments. Lower right: The DSGRN OR designs predicted to perform more robustly than the simple OR designs. Scoring: The difference in CDM scores between the low and high designs is shown in each row above the corresponding topology. Topology robustness scores predicted by DSGRN for the NOR and OR circuit topologies are shown in each row below the corresponding topology. These numerical scores should be interpreted ordinally rather than as absolute values with specific interpretation

The parts labeled with r# are associated to constitutively expressed CRISPR guide RNA (gRNA) gene products introduced in (40) that repress transcription when bound. Inducible versions of these parts were built in this study to use in tandem with the previously built constitutively expressed parts. Specifically, binding sites for *β*-estradiol and doxycycline hyclate were added to the gRNA promoters. In the presence of an inducer, the associated gRNA is expressed and represses the production of its downstream target, either another gRNA or green fluorescent protein (GFP).

The inducible and constitutively expressed gRNA parts are combined into circuits that act as sensors. For example, OR logic is realized when the absence of both inducers is associated to the absence of fluorescence while the presence of either inducer causes the production of GFP. In other words, the OR circuit acts as a sensor to signal the presence of one or both molecules.

### 1.3. Major contributions

The assessment of circuit performance was based on flow cytometry data. The flow cytometry samples frequently expressed bimodal distributions spanning the ON and OFF fluorescence states (Fig. 3). Given the lack of resources available to repeat the experiments, a machine learning method called Binary Event Prediction was developed to separate each bimodal distribution into an OFF distribution and ON distribution (Fig. 4). The sample was then classified as primarily ON or OFF depending on which distribution had greater mass. Using this technique useful information was extracted from suboptimal data.

**Fig. 3.**
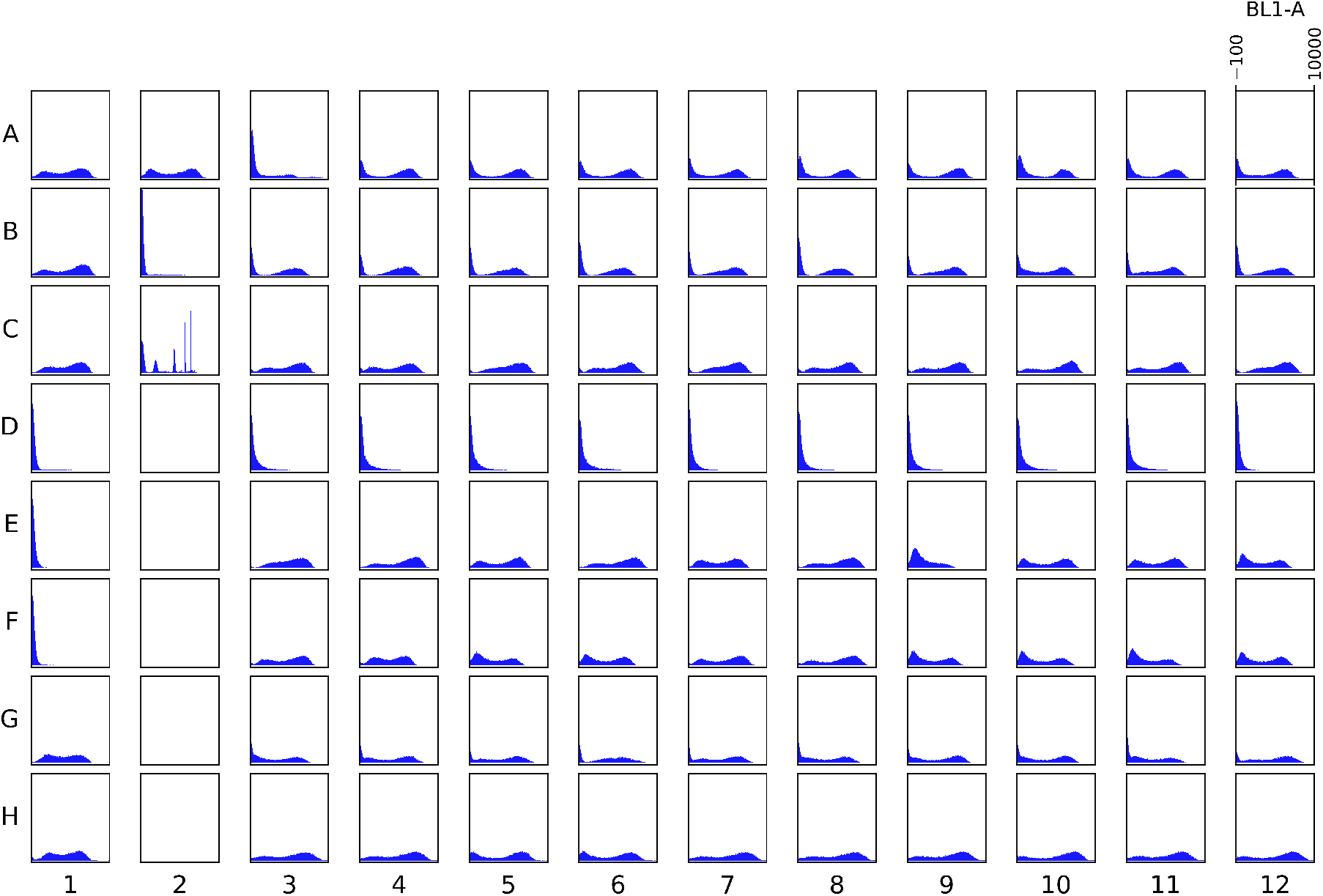
An example 96-well plate in SC media (plate id `r1fqsfkwxcccv6`, see Table 2) sampled 12 hours post-incubation showing fluorescence area distributions in arbitrary units. Controls (positive, negative, and bead) are in the first two columns. The remainder of the columns are data for the eight builds, one per row, with varying inducer concentrations.

**Fig. 4.**
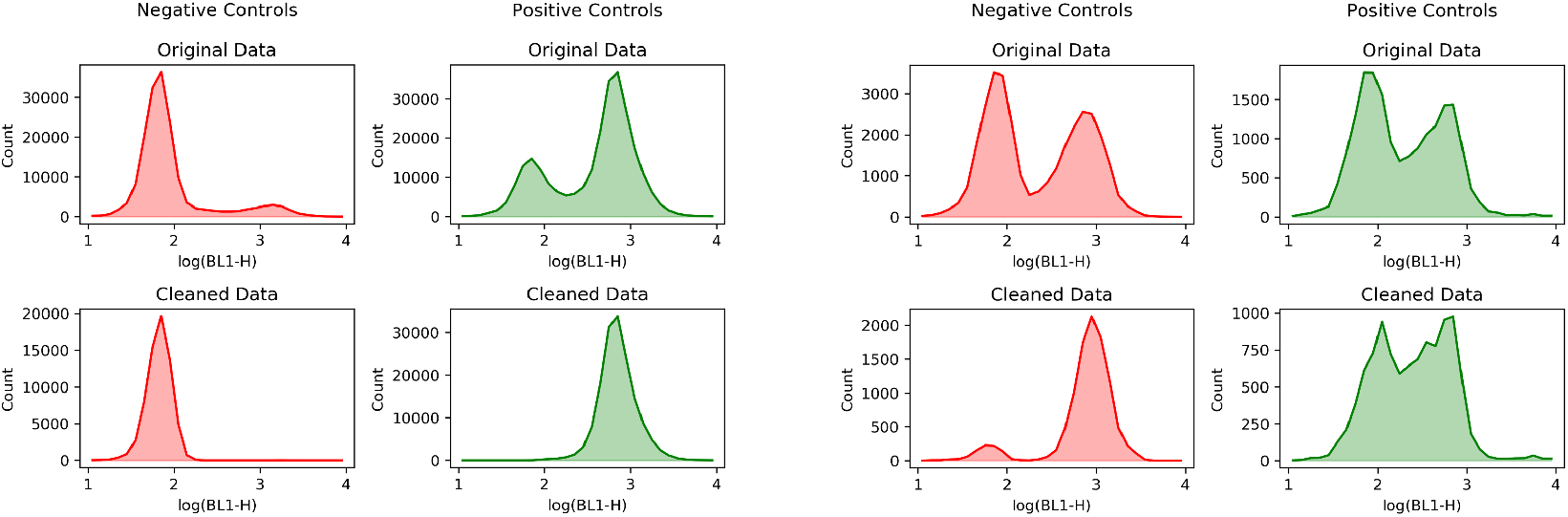
Examples of effective (left two columns) and ineffective (right two columns) cleaning of controls after stage one of Binary Event Prediction. Effective cleaning results in unimodal distributions, a low one for the negative control and a high one for the positive control. Ineffective cleaning either does not eliminate bimodality or does not result in the low (high) distribution being associated to the negative (positive) control. Negative controls are shown in red (first and third columns) and positive controls are shown in green (second and fourth columns). The histograms are fluorescence height measurements in log arbitrary units. Top row: Unmodified controls pooled over replicates. Bottom row: Cleaned controls pooled over replicates.

After separation of bimodality, the analysis showed that most of the builds responded with better performance than a null model (Fig. 5(a)-(b)). However, the performance of the engineered switches were not consistent with the CDM predictions of which parts would lead to high performing circuits and which parts would lead to low performing circuits (Fig. 5(f)), likely due to assumptions that were placed on the CDM method for this application (see Section 2.1.2). DSGRN predictions were partially fulfilled in that DSGRN NOR topology outperformed the simple NOR topology, but no appreciable difference existed between the DSGRN and simple OR topologies (Fig. 5(e)). There is therefore evidence that a more complex circuit may at times exhibit greater success across experimental conditions.

**Fig. 5.**
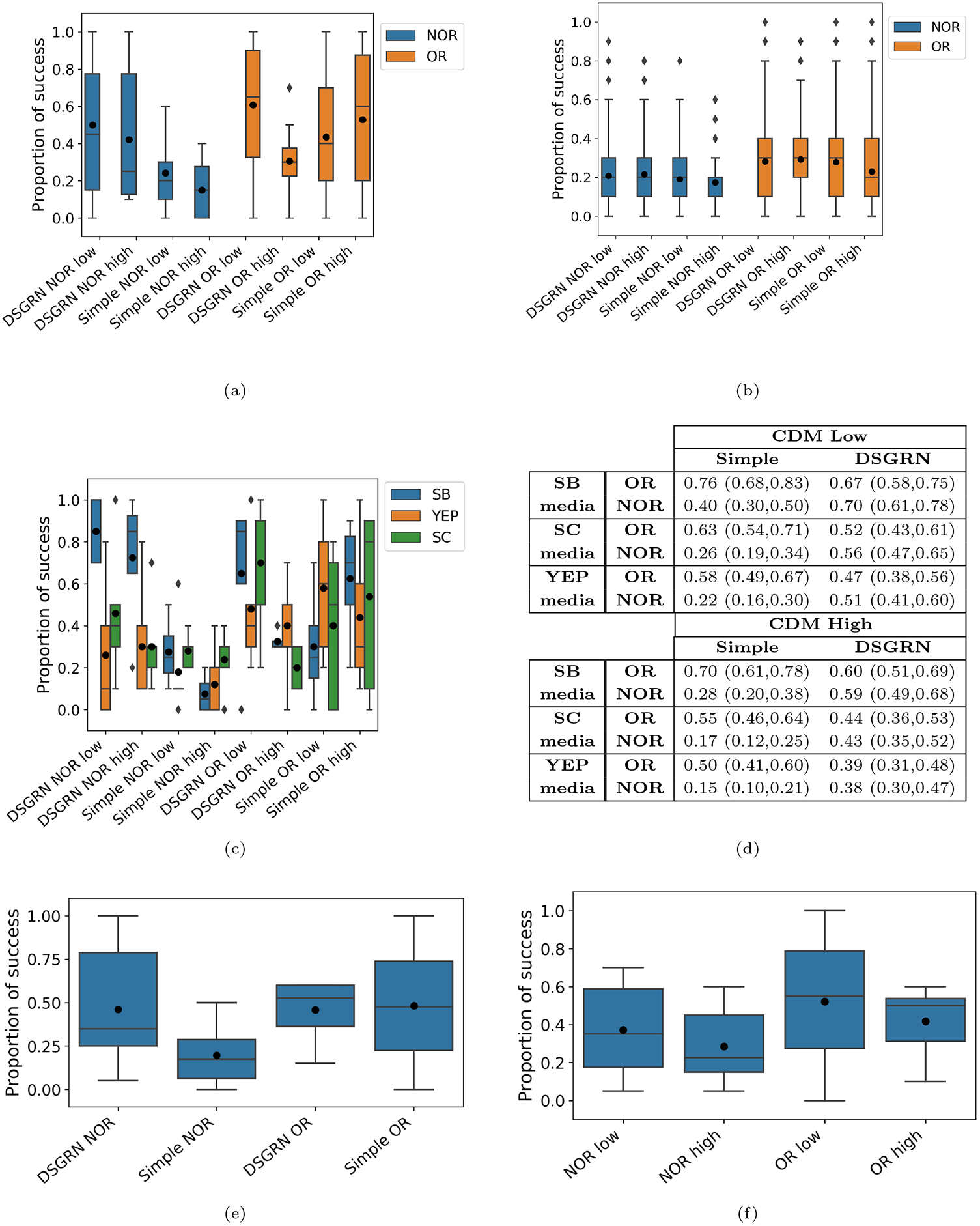
(a) Distributions of 14 plate performance scores (proportion of successes per build per plate) across the eight designs. The vertical bars indicate range with outliers as diamonds. The rectangles show 25-75 quartiles, with the horizontal bar showing the median and the black dot showing the mean. (b) Baseline distributions for plate performance scores; each plate had its BEP ratios permuted over all well and time points 1000 times. (c) Plate performance scores split by media (5 plates each for SC and YEP, 4 plates for SB). (d) Predicted probabilities of success for each build with 95% confidence intervals for the true probabilities of success. (e)-(f) Distributions of plate performance scores pooled across media and CDM design (e) to assess DSGRN predictions, and scores pooled across media and topology (f) to assess CDM predictions.

**Fig. 6.**
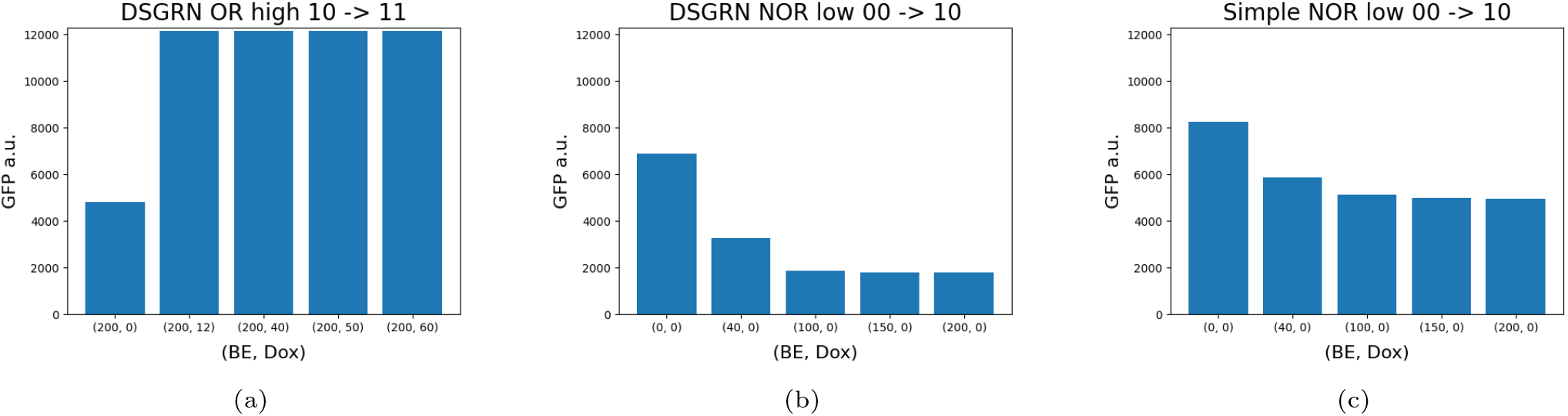
Example predicted steady state values of the geometric mean of the flow cytometry distribution of GFP a.u. from Hill function models. (a) The DSGRN OR/CDM high design as it transitions from an initial state of BE only to BE+Dox for various titration levels of Dox. The logical circuit should show a high fluorescence signal at all conditions. The BE-only steady state has a strong GFP signal in comparison to the low GFP steady state (1500-2000 a.u.), but is significantly lower than inducer conditions where Dox is present. (b) The DSGRN NOR/CDM low design as it transitions from an initial state of no inducers to BE at various titration levels. The condition (0,0) should show high fluorescence and all subsequent bars should show low fluorescence. (c) The same as (b) for the simple NOR/CDM low design. It is seen that BE does not effectively repress GFP signal in panel (c).

## 2. Results

### 2.1. Predictions

#### 2.1.1. DSGRN predictions of circuit topology performance

The DSGRN Design Interface provides robustness scores for a sample of network topologies exhibiting the desired truth table within a user-specified size range. Higher robustness scores correspond to predictions of greater success across experimental conditions. In principle, the robustness score can achieve a maximum of 1, but in reality this score is highly improbable. In Fig. 2, the DSGRN NOR topology has a score of 0.22, more than twice the simple NOR topology with a score of 0.08. Similarly, the DSGRN OR topology has a four-fold score of 0.28 compared to the original at 0.07. According to these predictions, the DSGRN topologies should exhibit better performance over condition space, which consists of media conditions and inducer concentrations in the experiments performed here. It is important to realize that DSGRN robustness scores are not to be taken as quantitative predictions. The robustness scores for the OR topologies would naively indicate that the DSGRN topology should successfully show the correct logic four times as often as the simple topology. However, that is at best an indicator of relative performance across all condition space (empirical numerical evidence for this can be found in (41)), not the limited condition space that is available experimentally. Therefore, only ordinal interpretation of the robustness scores is justified.

It is not always advantageous to build the circuit topology with the highest robustness score. As robustness increases, the redundancy in the network and therefore the complexity of the network is also increasing. There is a trade-off between predicted robustness and the ease of building the circuit. There were two NOR designs with the same number of parts as the DSGRN topology in Fig. 2 with a score of 0.30 that were rejected due to doubts about build feasibility.

#### 2.1.2. CDM predictions of various parts assignments

Given a circuit topology, it remains to assign parts to each node in the network. This was accomplished using a modification of the Combinatorial Design Model (CDM), a neural network model designed to use partial data to predict the performance of genetic constructs across untested experimental conditions. The data used to train the model were fourteen dosage response curves, a BE-inducible and Dox-inducible version for each of the seven gRNA promoter regions evaluated in this study: r1, r2, r3, r5, r7, r9, and r10.

CDM can be naturally used to predict the performance of simple NOR circuits; that is, it predicts the outcome of combining two inducible parts together when information is only known about individual parts. It is not explicitly designed to choose parts to construct larger circuits, and so assumptions had to be made in order to apply it. First, it was assumed that the independent CDM scores for NOR units could be combined in a way that leads to an accurate prediction of whole circuit behavior. Second, the model was trained on data for inducible parts only, but there were both inducible and constitutively-expressed parts in the parts library. It was assumed that the rank-ordering induced by CDM scores on combinations of inducible parts was preserved for the combinations of constitutively-expressed parts.

Using CDM predictions, two parts assignments were made for each circuit topology: one that was expected to be a better performer and another that was expected to be a poorer performer. These parts assignments are shown in Fig. 2, along with the difference between the high and low design CDM scores. While not interpretable in a quantitative manner, the difference in CDM scores are roughly comparable across circuit topologies. Again, only ordinal rankings are appropriate in the interpretation of these performance scores. As an example, compare the 1.49 difference in low and high simple NOR designs to the 2.54 difference in simple OR designs. The interpretation is that a larger difference between low and high designs should be seen in the simple OR topology versus the simple NOR topology. As a side effect of parts assignment, CDM was also able to reduce the intervals of inducer concentrations to be tested.

### 2.2. Data Overview

There is a minimal amount of information the reader needs about the experiments to interpret the following two sections that contain the major contributions. The eight builds in Fig. 2 were grown in three media types, YEP 2% Dextrose (rich media), Synthetic Complete (standard media), and Synthetic Complete containing 1% Sorbitol (slow growth media). The three media will be referred to as YEP, SC, and SB, respectively. The cultures were used to create fifteen 96-well plates, with five plates per media condition. Each plate had titrated inducer concentrations for one or both of *β*-estradiol (BE) and doxycycline hyclate (Dox), see Appendix Table 1. The five titration experiments were BE titration into base media, Dox titration into base media, BE and Dox simultaneous titration into base media, Dox titration into BE spike-in media, and BE titration into Dox spike-in media (the term spike-in means that the inducer was added to the media before the incubation period). Flow cytometry (FC) data were collected at three time points: 12, 24, and 36 hours after an incubation period of 16 hours. An event refers to a single flow cytometry measurement and a sample is the collection of events associated to a well (a fixed location on a plate) at a single time point. The presence/absence of the inducers determines the expected outcome of the GFP signal in the FC measurements for the OR and NOR circuits; see Appendix Table 2 for a key to the media and expected outcomes of the circuits for each plate.

**Table 1.** The inducer concentrations in nM for each logical transition. For each plate there is a distinct logical transition that represents the change between the replicate columns 3-4 on each plate in comparison to all subsequent plate columns (recall that plate columns 1-2 are controls). Each logical transition corresponds to three plates, one for each media condition (see Table 2), and each inducer concentration was sampled at three time points during the experiment. To read the table, consider any pair of inducer concentrations (B,D). The Boolean translation of this pair is B=0 if BE is absent and B=1 otherwise; similarly, D=0 if Dox is absent and D=1 otherwise. So the first row 00 → 10 indicates that there are no inducers in the “Plate cols. 3-4” column, but that BE is present at some titration level in all the rest of the columns. This combination of Boolean states for BE and Dox allows the assessment of two different circuit input states for each design on each plate at each time point. The last two rows 01 →11 and 10 →11 represent the spike-in conditions, since Dox or BE is present in the inducer pair in “Plate cols. 3-4”.

| Logical input transition | Inducer concentrations per plate: (BE, Dox) in nM |  |  |  |  |
| --- | --- | --- | --- | --- | --- |
|  | Plate cols. 3-4 | Plate cols. 5-6 | Plate cols. 7-8 | Plate cols. 9-10 | Plate cols. 11-12 |
| 00 → 10 | (0, 0) | (40, 0) | (100, 0) | (150, 0) | (200, 0) |
| 00 → 01 | (0, 0) | (0, 12) | (0, 40) | (0, 50) | (0, 60) |
| 00 → 11 | (0, 0) | (42.5, 12.25) | (85, 24.5) | (170, 49) | (212.5, 61.25) |
| 01 → 11 | (0, 40) | (40, 40) | (100, 40) | (150, 40) | (200, 40) |
| 10 → 11 | (200, 0) | (200, 12) | (200, 40) | (200, 50) | (200, 60) |

**Table 2.**
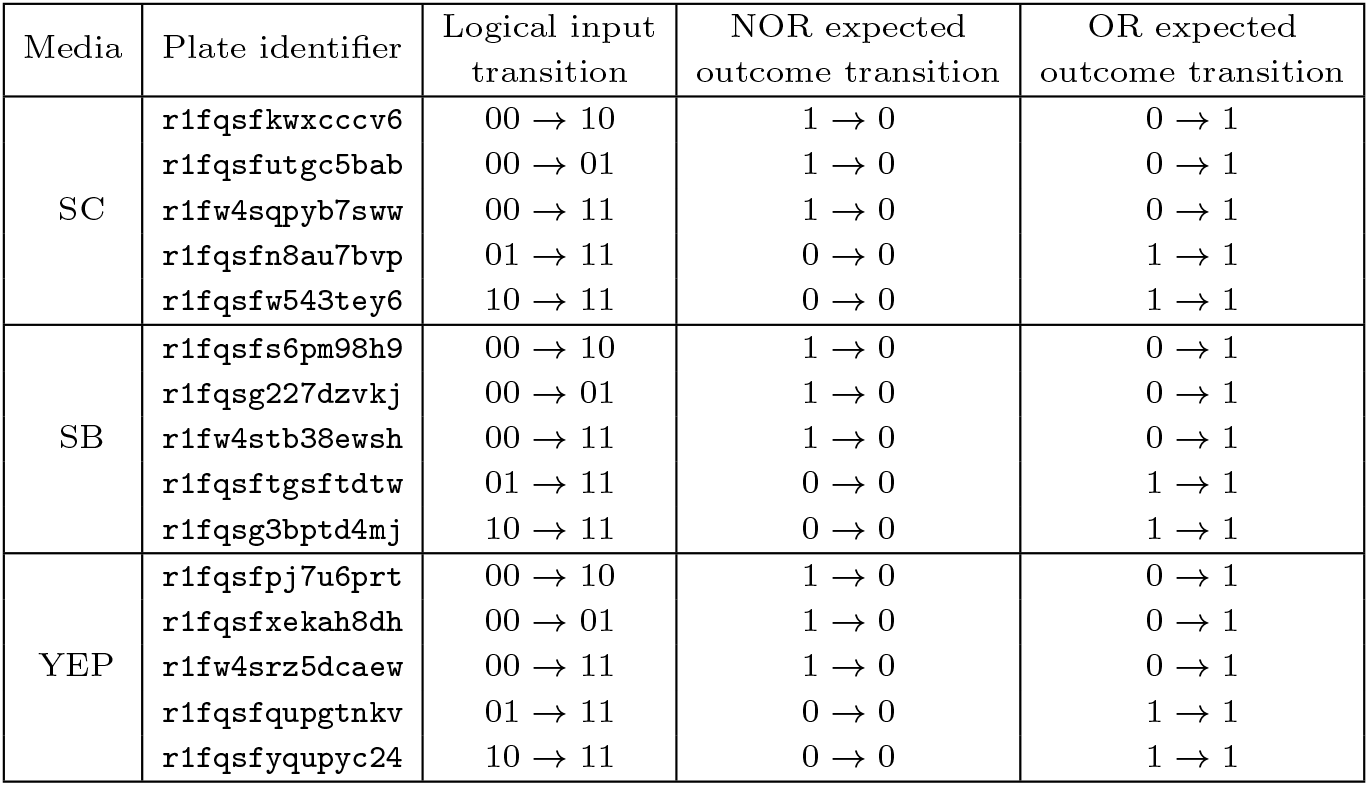
Media condition and logical transitions for each plate’s unique identifier. For the logical input transitions, BE is in the first place and Dox is in the second place; see Table 1 for details. The expected outcome transition indicates whether GFP is supposed to be produced or not. For NOR, case 0 → 0 means that the NOR circuit should not be producing GFP in either input transition condition, while case 1 → 0 means that the NOR circuit should stop producing GFP after an initial growth period where it has been active. Similarly, OR is tested on the outcome transitions 1 → 1 and 0 → 1. The success of the outcome transition is dependent on the temporal processes of induction and GFP decay.

### 2.3. Data Processing

The RT provided standardized FC data products for visualization and analysis. Plots of these data showed significant bimodal distributions in both the OR and NOR builds, see Fig. 3 for an example plate. A possible explanation for bimodality in the data is that induction times for the circuits are slow, and therefore there are populations of both sufficiently and insufficiently induced cells.

However, there is also bimodality in the positive and negative controls, see the first column in Fig. 3 as well as the first row of Fig. 4. In the positive controls, this suggests potential mutations where the GFP coding region has been compromised. In the negative controls, this suggests possible contamination of the plates with GFP producing cells. Given these issues with the controls, it would be ideal to redo the experiments.

In many situations, it is not feasible to redo experiments either due to lack of resources or inability to obtain more samples. This is the case for the data presented here; another set of experiments was not possible to obtain. The challenge was to see how much information could be drawn from data with anomalous controls. One solution would be to presume that the GFP-positive population in the negative control is uniform throughout the plate and subtract that portion of the population from all of the wells. This technique has the significant disadvantages that (i) the assumption of uniform contamination may be incorrect, (ii) it takes significant manual effort, and (iii) it uses only one channel out of 16 to judge the identity of problematic events.

To address these issues, a machine learning technique referred to as Binary Event Prediction (BEP; (42)) was developed to separate every bimodal distribution into low and high (or OFF and ON) distributions on a per-plate basis using 16 flow cytometry channels, each with width, height, and area measurements. BEP uses a random forest classifier in two stages. In the first stage, the pooled replicates of the positive and negative controls on a plate are used to train a random forest classifier. These classifiers, one per plate, are then used to predict the training data. Those events that exceed an adaptive probability threshold (see Methods 4.5.2) for either the high distribution in the positive control or a low distribution in the negative control were kept, producing “clean” controls that often achieve reduced bimodality; see Fig. 4 for examples of effective versus ineffective cleaning. In stage two of BEP, the cleaned controls trained new random forest classifier models, one per plate, that were used to predict the ON/OFF status of all events on the plate. Appendix Fig. 8 shows the fluorescence distributions for the original and cleaned controls for each of the 15 plates in the experiments. On the left are 9 plates with effective cleaning of the controls, i.e. they have achieved unimodality after cleaning, and on the right are the remaining 6 plates that show ineffective cleaning.

**Fig. 7.**
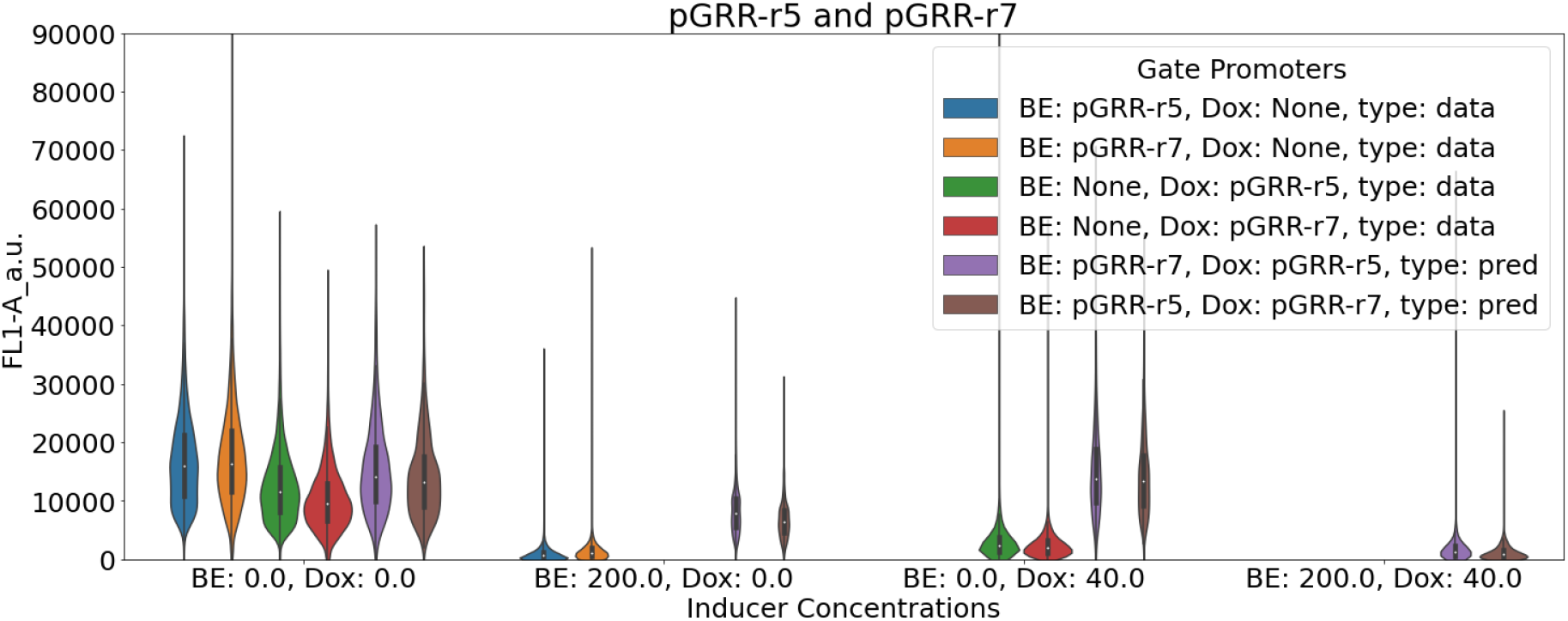
Example CDM prediction based on parts r5 and r7. The *y*-axis shows fluorescence in arbitrary units, and the *x*-axis has combinations of the inducers *β*-estradiol (BE) and doxycycline-hyclate (Dox). Blue and orange denote data taken for parts r5 and r7 respectively in the presence of a high concentration of BE. Likewise, green and red show data for r5 and r7 respectively in the presence of a high concentration of Dox. Purple and brown show CDM predictions for the combination of BE-inducible r7 and Dox-inducible r5 and for the combination of BE-inducible r5 and Dox-inducible r7, respectively, at all inducer conditions, including the case where both BE and Dox are present and no data were gathered.

**Fig. 8.**
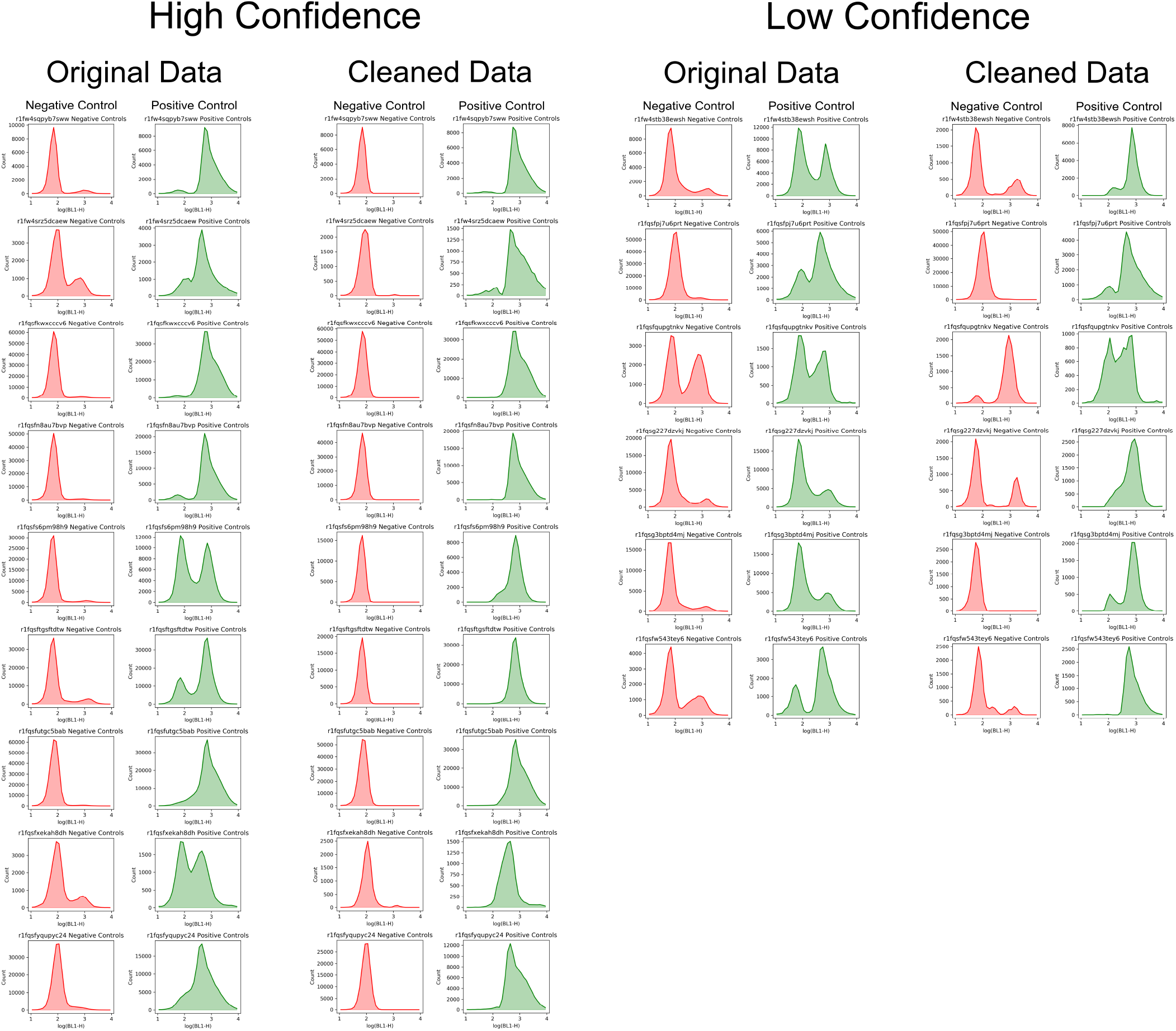
This figure shows the GFP height distribution for the negative and positive controls pooled across replicates for each of the 15 plates, including the dropped plate `r1fw4stb38ewsh` shown in the first row in the right two columns. The 9 plates on the left had an effective cleaning procedure, resulting in unimodal low (high) distributions for the negative (positive) controls. The plates on the right did not achieve unimodality in one or both controls.

A comparison was performed between the original log-transformed data and a subset of the data identified by BEP in order to assess BEP performance. The BEP subset for each sample was determined by comparing the number of BEP-predicted ON events versus predicted OFF events. Whichever predicted distribution comprised more than 50% of the total events was used as the modified sample data. Both the original and modified datasets were analyzed via the Round Trip tools (i) Performance Metrics (PM; Methods 4.5.3) to assess fold change between expected OFF and expected ON circuit states and (ii) Data Diagnosis (DD; Methods 4.5.4) to assess variability in the fold change as function of experimental variables. PM computes the fold change in median between all samples that are expected to be ON according to circuit function and all of those that are expected to be OFF. The distributions of the median fold change per plate for the original and modified datasets are shown in Appendix Fig. 10. The distributions are seen to be very similar, indicating that in aggregate the fold changes between circuit ON and circuit OFF states are not improved by the BEP technology. However, DD shows that even though the fold change hovers around 1 for both datasets, the BEP subset data has many more outliers at higher fold changes (*≥* 5) than the original data; compare Appendix Figs. 11 and 12. This indicates that circuit performance measured as fold change is greatly improved for a number of samples.

**Fig. 9.**
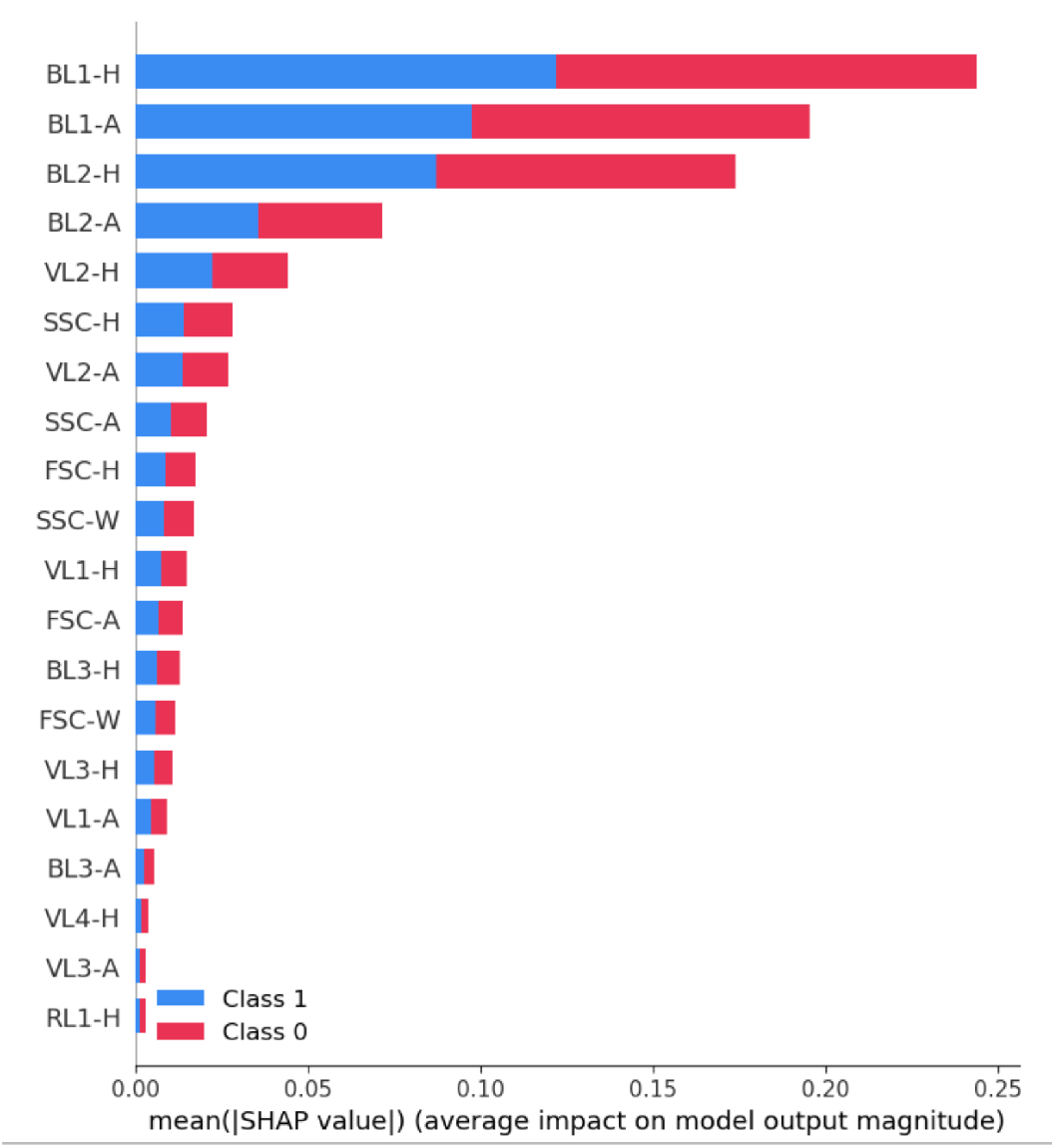
SHAP values showing the importance of various flow cytometry channels to the BEP classifier. The primary GFP channel is BL1, with the -H,-W, and -A modifiers indicating height, width, and area, respectively. BL2 is the secondary GFP channel. FSC and SSC are forward and side-scatter, respectively, that roughly correlate with cell size and shape. FSC and SSC are the only channels that measure characteristics other than fluorescence.

**Fig. 10.**
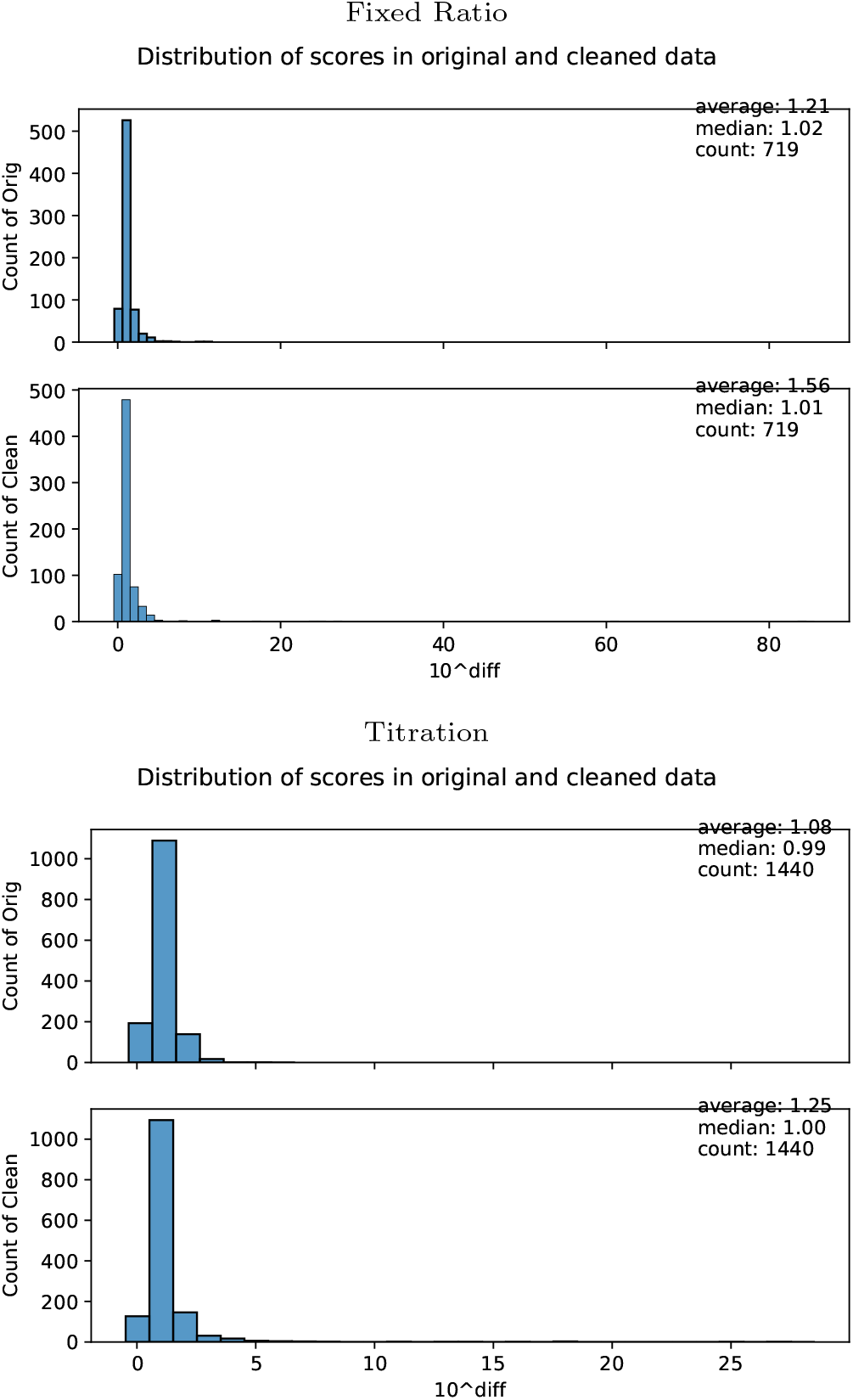
Fold change distributions for the clean model and the unmodified log-transformed data computed by Performance Metrics. Top: Three plates in the dual-inducer experiment. Bottom: Twelve plates in the single-inducer experiment.

**Fig. 11.**
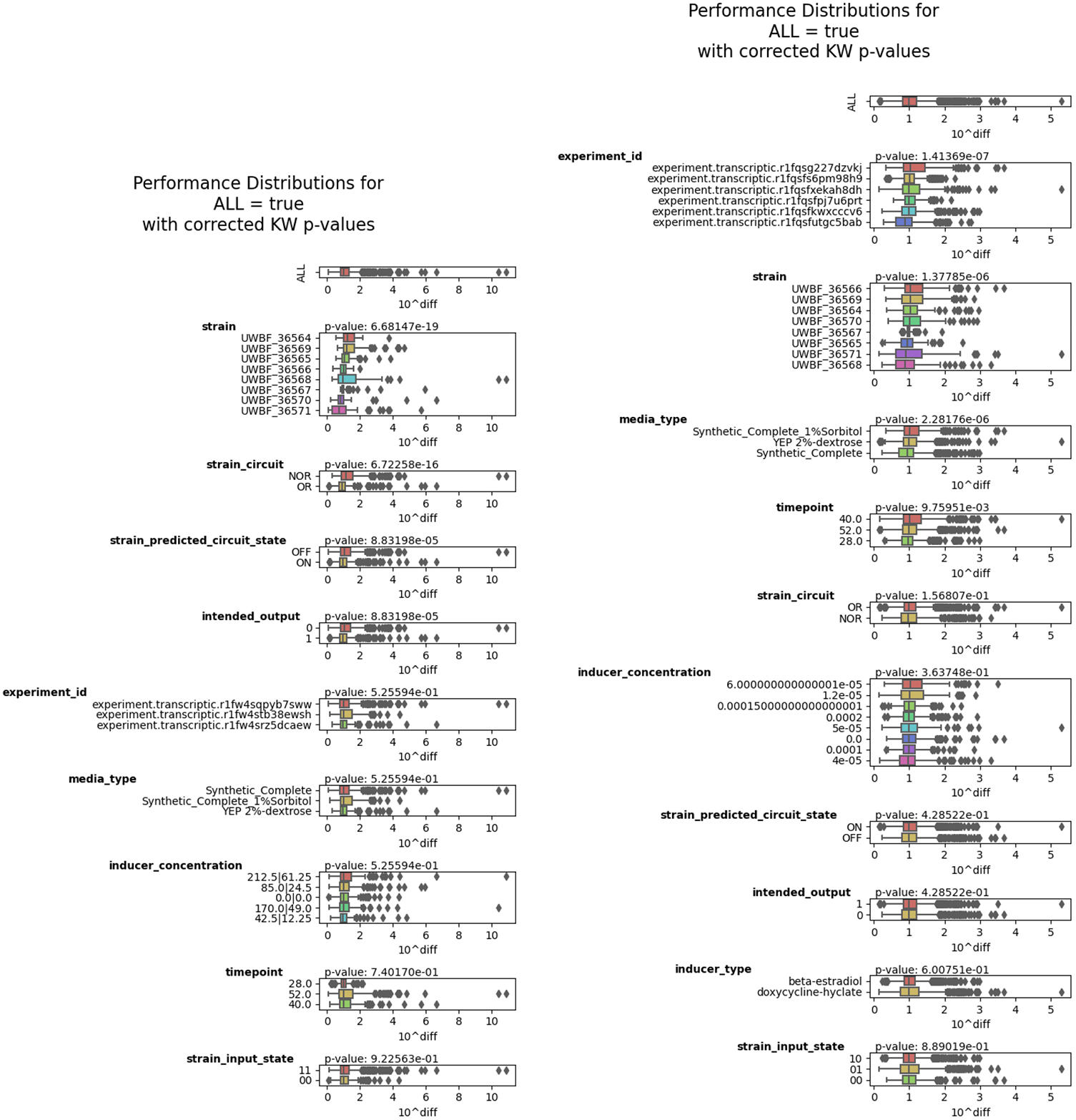
Data Diagnosis analysis of original log-transformed data. Left: Three plates from the dual inducer titration experiment. Right: Twelve plates from the single inducer titration experiment. The plate identifier is given under the label `experiment_id`. See Table 2 for information on the plate conditions. The plate dropped from further analysis is `r1fw4stb38ewsh`, one of the three plates on the left.

**Fig. 12.**
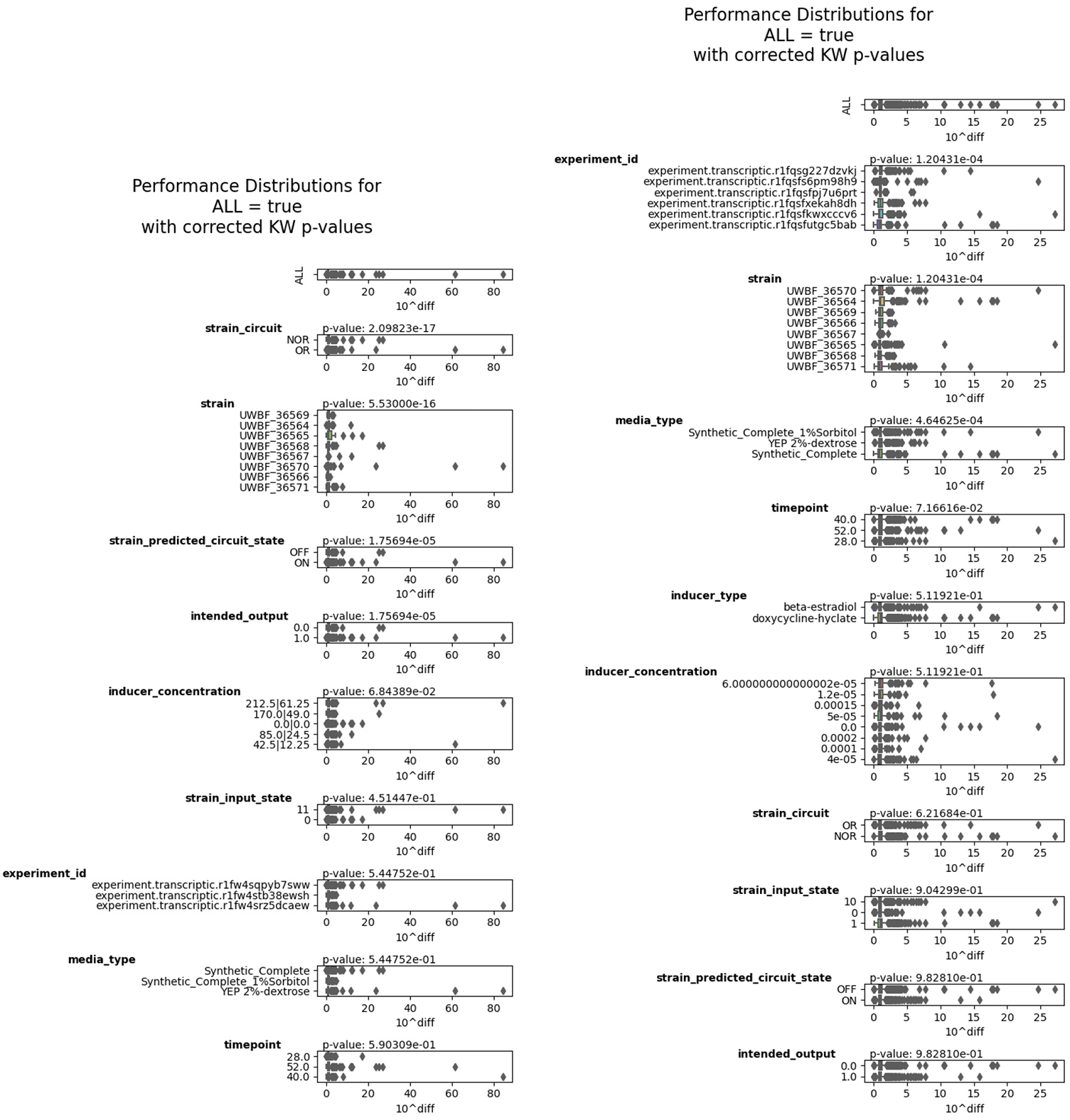
Data Diagnosis analysis of clean model showing the presence of large outliers in comparison to the raw log10-transformed data in Fig. 11. Left: Three plates from the dual inducer titration experiment. Right: Twelve plates from the single inducer titration experiment. The plate identifier is given under the label `experiment_id`. See Table 2 for information on the plate conditions. The plate dropped from further analysis is `r1fw4stb38ewsh`, one of the three plates on the left.

The BEP methodology reduces human labor in sorting through data and draws conclusions using all 16 channels and all 3 time points of the flow cytometry data without assuming a uniform distribution of contamination. As is intuitive, the GFP channel was the most important contributor to the model according to SHAP values (43), see Appendix Fig. 9. Moreover, as will be demonstrated in the next section, the BEP event predictions of ON versus OFF state permit the evaluation of circuit performance, which is otherwise problematic when the fold change between expected ON and OFF distributions is approximately 1.

### 2.4. Data Analysis

Circuit performance was assessed by examining the three time point flow cytometry distributions for GFP for each well in a plate. Each FC distribution was required to meet conservative criteria for cell presence in order for a well to be assessed for success. The total number of events in the distribution had to be at least ten thousand and there had to be a cell density of at least 500,000 cells/mL. If any time point did not meet these criteria, then the well was classified as a failure, with the following exception: one plate (`r1fw4stb38ewsh`, see Appendix Table 2) with SB media had such wide-spread insufficiency in cell density and number of events that the whole plate was excluded from analysis. This plate was one of the plates with ineffectively cleaned controls (see first row, right columns in Appendix Fig. 8) and was an SB media plate. The remaining fourteen plates consist of five plates for each of SC and YEP media and four with SB.

Successful performance was assessed by the following scoring technique. For each time point, there is a proportion of events in the FC distribution that are predicted by BEP to be ON (*p*) or OFF (1 *− p*), where *p* will be called the BEP ratio. If for a well all three time points satisfied *p >* 0.6 (*p <* 0.4), then the build was declared to be exhibiting ON (OFF) behavior. Additionally, if neither condition above was met but *p* increased (decreased) over time, then the build was said to exhibit ON (OFF) behavior. This latter condition was chosen since some of the builds may still have been undergoing the temporal processes of induction and GFP decay over the course of the experiment. Any well that did not meet one of these criteria was said to have indeterminate behavior and was classified as a failure. A success for a well is defined to be any case where the exhibited behavior matched the intended ON or OFF state of the circuit given the presence or absence of the inducers, otherwise it was classified as a failure.

Successful or unsuccessful performance could be characterized by looking at the distribution of well successes across all plates for each build. However, since BEP exhibited variable performance on a per-plate basis, successes were aggregated on the plate level for each build. A plate performance score for a build is the number of successes divided by ten, since each build was measured under five inducer conditions with two replicates on each plate.

The fourteen plate performance scores form the distributions plotted in Fig. 5(a). The category labels indicate the topology, circuit, and CDM prediction in order. Median performance shows a wide range of values and the interquartile ranges are generally large, therefore it is unclear whether or not any of these circuits can be declared successful on the whole. To address this, a baseline distribution was created where each plate had its BEP ratios permuted over all 8 builds and 3 time points 1000 times. The baseline plate performance score was computed for each circuit at each iteration and the distributions are shown in Fig. 5(b). Although there is overlap in the interquartile differences, the means and medians of the true scores for DSGRN NOR, DSGRN OR/CDM low designs, and simple OR topologies are all higher than those of the randomized empirical null.

In Fig. 5(c), the distributions in Fig. 5(a) are split over media condition. The distributions only have 4-5 points each, but the boxplot shows suggestive patterns. First, the DSGRN NOR topologies perform either better or similarly to the simple NOR topologies in all three media conditions, with particularly high performance in the SB media condition. Second, the underperformance of the DSGRN OR/CDM high design seems to be due primarily to the SC media. Disappointingly, no build seems to exhibit similar, and therefore reproducible, performance across media conditions. The closest is the simple OR/CDM high design.

To provide some statistical quantification of the observed trends, the probability that a new observation will be a success for a given build and media condition was estimated using a logistic regression model (Appendix D). Fig. 5(d) shows these predicted probabilities along with a 95% confidence interval for the true probability of success. When the confidence intervals do not overlap, that is suggestive of, though not a guarantee of, significantly different probabilities of success. Given a media condition and circuit, the only time there is non-overlapping confidence intervals, and therefore possible significance, is between the simple NOR topologies with lower predicted success and DSGRN NOR topologies with higher predicted success. This can be seen by choosing a NOR circuit row and pairwise comparing the confidence intervals between the simple and DSGRN topologies. Additionally, by examining the top half and bottom half of the table, it can be seen that the CDM low designs have higher probabilities of success than the CDM high designs, but there is overlap in the confidence intervals.

The accuracy of the DSGRN robustness predictions and the CDM performance predictions are further assessed by pooling media conditions and builds together, as shown in Fig. 5(e)-(f). In Fig. 5(e), the plate performance scores for the high and low CDM builds are pooled to compare the DSGRN topologies against the simple topologies. The DSGRN NOR topology outperforms the simple NOR topology, indicating that that circuit redundancy may indeed produce more robust performance, as also supported by the statistical analysis in Fig. 5(d). On the other hand, the two OR topologies are not substantially different performers (in mean and median). But Fig. 5(a) and (d) show that the DSGRN OR designs show a large differential performance from each other, whereas the two simple OR designs do not. A possible contrary conclusion is that the DSGRN OR topologies are not as reproducible as the simple OR designs in terms of the specific biological parts deployed, which is counter to DSGRN predictions. In Fig. 5(f), the plate performance scores for the DSGRN and simple topologies are pooled to compare the CDM low performance predictions to the CDM high performance predictions. The CDM low designs show a trend of higher performance than CDM high designs, consistent with Fig 5(d).

The simple NOR design is an anomalously poor performer compared to all other designs. This is interesting because all the other builds contain similar NOR gates, although they use different gRNA parts (see Fig. 2). Either the simple NOR builds are unexpectedly fragile, or there is perhaps synergistic activity when multiple NOR gates act in concert. To further explore this performance failure, a Hill function ordinary differential equation model of the designs was created using parameter fits (Methods Section 4.5.5) from the same dosage response experiments used to train CDM.

In general, the Hill model predicted that circuits should respond more strongly to Dox (Fig. 6(a)), but that the dosage response to BE was acceptable (Fig. 6(b)) except for the two simple NOR designs, in which the presence of BE alone is predicted to fail to turn the circuit OFF (Fig. 6(c)). Fig. 6(a) shows the DSGRN OR/CDM high design and illustrates the difference in BE and Dox performance. A success is a high GFP signal in all five bars, where the left-most bar is BE alone at its highest titration concentration and the remaining four bars are combinations of BE and Dox, with Dox at various titration levels. High GFP signal is achieved even for the BE-alone condition, since the low GFP steady state corresponded to about 1500-2000 a.u., substantially lower than the left-most bar. However, the BE-induced GFP signal is markedly lower than that for the BE+Dox combinations. Dox in isolation produced GFP signal in comparable amounts to BE+Dox (Appendix E).

Fig. 6(b)-(c) shows the differential Hill-model predicted performance of the DSGRN and simple NOR topologies using parameters for the CDM low designs. In this case, BE successfully represses the signal of the DSGRN NOR design to the 1500-2000 a.u. low steady state, but does not suppress that of the simple NOR design. In the absence of both inducers (first bar) both designs are predicted to exhibit a high fluorescence signal, but in the subsequent titrations, the simple design is predicted to have a GFP signal more than twice the low steady state. The BE-inducible parts r10 and r5 are used only in the simple NOR designs and not in the other designs (Fig. 2). One cannot exclude the hypothesis that the inclusion of the BE-inducible parts r5 and r10 into the three other topologies would also result in degraded performance.

## 3. Discussion

This manuscript introduces a set of tools for the design of circuit topology through performance prediction and a set of tools to improve the efficiency and accuracy of building synthetic circuits. Together this Design Assemble toolchain is melded into a previously published toolchain, the Round Trip. During the DA portion of this sequence, OR and NOR logical circuit topologies and parts assignment were designed using the predictive tools DSGRN (29; 38) and CDM (32) and built using the laboratory software tools DASi (33; 34), Terrarium (36), and Aquarium (37). The Round Trip (25) portion of the toolchain enhanced the reproducibility of experimental results through the automation of experimental specification, subsequent data handling, and standardized analysis.

During data inspection, it became clear that contamination, of the negative controls with GFP-producing cells and of the positive controls with mutated cells lacking GFP production, was a real concern—appearing as strong bimodality in the FC distributions. However, there were insufficient resources to repeat this suite of experiments, and no guarantee that the issues would be resolved through replicated experiments. A challenging goal is the development of methods that can use all data, whether optimal or suboptimal, to inform future experiments. To this end, a machine learning technique called Binary Event Prediction (BEP) (42) was developed to separate bimodal distributions based on 16 flow cytometry channels. The intent was to identify true ON/OFF versus false ON/OFF events in order to assess circuit performance. Aside from this, BEP provided extra information that flow cytometry channels correlated with cell size were not important in the classification.

Although the simplest topologies for OR and NOR circuits are well-known to synthetic biologists, there are many circuit topologies that also exhibit the correct logical behavior. DSGRN predicts that many OR and NOR circuit designs with redundancy should show more robust behavior over experimental conditions. CDM predicted groupings of parts that should show better versus worse circuit performance.

The simple and DSGRN NOR circuit performance exhibits a trend consistent with DSGRN predictions of circuit robustness, namely that a circuit design with redundancy may indeed be more robust with regard to diverse experimental conditions. However, there are mixed conclusions to be drawn for the OR circuit. On the one hand, DSGRN OR topologies show a smaller interquartile range, consistent with greater average reproducibility. On the other hand, the CDM-predicted high and low builds of the DSGRN OR topology show a much greater difference in median than the analogous builds for the simple OR designs, indicating lower reproducibility.

The designs which CDM predicted would be high performance were often outperformed by those CDM predicted would be low performers. This is likely due to faulty assumptions when creating a purpose-built version of CDM for this application. Either the scores for combinations of NOR genetic subunits were incorrectly calibrated or the assumption that genetic parts with gRNA binding sites exhibited similar behavior to their BE and Dox-inducible counterparts was incorrect. An alternative future route is to combine DSGRN topology generation with stochastic parts assignment optimization, such as that in Cello (15), or with the Wasserstein metric-based parts optimization in (17).

The trends in performance are suggestive rather than significant. These trends can be used to hypothesize the most promising directions for future experimentation. Possible avenues of exploration involve the choice of topology, particularly for the OR circuit, the choice of parts assignment, particularly for the simple NOR designs, the choice of experimental conditions, or the experimental platforms and protocols used. From the work here, better experimental controls and different assignments of parts are clear choices. A wider range of parts assignments and experimental conditions would lead to a better exploration of the operational envelope of the construct, and would therefore put higher confidence in the topology rankings produced by DSGRN. Lastly, alternative topologies for OR logic could provide further evidence for or against increased robustness of performance when comparing structural redundancy to build complexity.

In conclusion, synthetic logic circuits predicted to exhibit OR and NOR logic behavior were designed, built, observed using flow cytometry, and analyzed for performance using a toolchain designed to enhance robustness and reproducibility. A machine learning technique was developed and trained on all measured flow cytometry channels to mitigate unexpected experimental complications. The results indicate that more complex circuits exhibiting structural redundancy may provide more robust and reproducible behavior in a synthetic biology setting despite increased build difficulty.

## 4. Methods

### 4.1. Design tools

#### 4.1.1. DSGRN design software

Dynamic Signatures Generated by Regulatory Networks (DSGRN) is a Python package (29) that permits a user to comprehensively describe the long-term dynamical behavior of a network topology. In this context, it predicts the equilibrium behavior of a logic circuit under DSGRN parameterizations. A DSGRN parameterization is actually a large region of high-dimensional parameter space across which equilibrium behavior is constant. This is a finite decomposition of high-dimensional parameter space, called the DSGRN parameter graph, so that DSGRN exhaustively predicts all behaviors available to a network topology. To predict the robustness of a dynamical behavior, DSGRN computes the percentage of the finite number of regions in the DSGRN parameter graph that exhibit the behavior of interest.

While powerful and flexible, DSGRN currently requires custom scripting to ask specific questions, such as the enumeration of the range of behaviors of a synthetic biology construct. A wrapper Python package `dsgrn_design_interface` (38) was written to permit a plain language interface for a non-expert user to design a collection of feed-forward network topologies that satisfy a desired logic circuit behavior. Using this tool, thousands of network topologies can be screened for the desired logic behavior.

The input is a json configuration file, where the user specifies the desired circuit, whether the inducible inputs are activated or repressed, the size of the network topology, and the local logic of individual parts that are available to build the circuit (i.e. independent or dependent repressors or activators). The output is a collection of files accessible through an automatically generated Jupyter notebook that can be browsed for final choices of network topologies. An SBOL2 (Synthetic Biology Open Language) (28) formatted document can be created for any of the desired topologies. Comprehensive documentation is provided.

Given the user inputs, a DSGRN design parameter or set of parameters is identified that is consistent with the local logic of the parts. A successful output topology is one that exhibits the desired logic behavior at a design parameter. The neighboring parameters to the design parameters in the DSGRN parameter graph are well-defined, and are associated to small malfunctions in the intended operation of the circuit. Each successful output topology is accompanied by a numerical score that rank orders networks according to their predicted robustness based on these neighbors. For some successful network, let *D* be the number of design parameters with the correct equilibrium behavior, let *N* be the number of neighbors of all design parameters, let *N*_*c*_ be the number of these neighbors that exhibit the correct logic, and let *E* be the number of essential DSGRN parameters (44). Then the topology robustness score (e.g. Fig. 2) assigned to the network is a value between 0 and 1 defined by

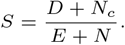

See (30) for a similar scoring method in another design application of DSGRN.

#### 4.1.2. Combinatorial Design Model (CDM)

Experiment and circuit designs are often additive, namely, single conditions are combined to elicit a collective response. For a circuit constructed of multiple parts, the challenge is to select the best combination of parts that produces the best response (e.g. the largest dynamic range). While there are tools that exist to support this task, such as Cello (16), they require a thorough characterization of every part (e.g. a titration of inducer for each part) to construct the circuit. The CDM is a neural network based model that is trained with a subset of conditions and predicts the left out conditions. The conditions need not cover a full range of titrations, but the predictions will then be limited to only the combinations of inducer concentrations and gates tested. For this effort, ON/OFF conditions for each part were tested and the flow cytometry distributions were predicted over all combinations of inducers and parts (e.g. Fig. 7). The CDM was structured to make predictions using multiple stages, each stage aligned with a given topology. For example, the six-node DSGRN NOR topology with two input nodes, an intermediate node, and two nodes with promoters at the GFP coding region had a three stage model corresponding to each of these “layers”. At each layer, a measure of deviation of predicted fold change was computed, with details of normalization varying at each stage. Since CDM scores change with topology size, it is more comparable to look at differences between high and low scores as in Fig. 2, rather than to interpret absolute scores.

### 4.2. Build tools

#### 4.2.1. Aquarium

For plasmid and yeast construction, Aquarium software (14; 37) was used to manage inventory, experimental workflows, and protocol execution. The input to Aquarium is a workflow, or linked set of protocols. During the execution of a workflow, data and metadata are logged and tracked to a database, which is used for future automated planning. Aquarium workflows are executed by technicians following on-screen instructions generated by Aquarium. Occasionally, the same protocol is executed by multiple technicians at the same time or sequentially (one technician leaving and another starting where the other left off). Technicians are trained beforehand, but had various skill levels and experience. During execution of the protocols for the build steps and parts library dosage response experiments conducted at the UW BIOFAB (45), technicians were unaware of project goals and were instructed to exactly follow the steps and instructions provided by Aquarium.

#### 4.2.2. Terrarium

There are two inputs to the computer-assisted process planning software Terrarium (36): metadata from previously run Aquarium workflows and a biological design encoded in a biological manufacturing file (BMF) using the JSON (JavaScript Object Notation) standard. A BMF defines three components: (i) the laboratory configuration, (ii) relationships between biological sub-components in a given design, and (iii) biological definitions, which include the type of biological sample (yeast, bacteria, DNA, sequences, etc.). Terrarium uses lab inventory information from the BMF plus metadata from previously executed Aquarium workflows to generate a model of the laboratory. This enables predictions for lead time, costs, labor, and predictions of experimental errors for proposed workflows that permit the creation of an optimized Aquarium workflow.

#### 4.2.3. DASi

In some cases the sequence provided in a BMF is not available in the laboratory as inventory. In these cases, DNA sequences can be created in the laboratory by a combination of DNA subcloning or DNA synthesis from outside vendors. For this type of planning, Terrarium uses a planning subroutine, DASi (33), to automatically generate an economical DNA cloning assembly plan from sequences, using available lab inventory whenever possible. DASi has flexible input requirements for a DNA sequence, and may be either a string of characters, a GenBank file, an SBOL file, or a BMF. Unlike other cloning automation software, DASi does not require further design specifications beyond the DNA sequence to effectively create DNA assembly plans. Once an assembly is returned from DASi, the output file can be converted into a BMF that is processed by Terrarium into a validated Aquarium workflow.

### 4.3. Round Trip (RT)

DART relies upon the RT tool chain (25) to test assembled designs and then process experimental data. The RT accomplishes this by accepting a semi-structured Experimental Request (ER). The ER represents human-readable descriptions of the experiment and a set of structured tables that describe controls, builds, reagents, conditions, and measurements. The RT uses a tool called the Intent Parser (46) to resolve human-readable terms to their representations in SBOL, including their design and assembly. The Intent Parser validates the ER and converts the experiment description into a machine-readable format that goes through experimental planning. The experimental planner, XPlan (13), partitions and allocates the requested builds, conditions, and measurements to experimental runs that are launched at a laboratory with compatible software.

The RT processes the data through three main steps: a) database ingest, b) standardization and versioning, and c) automated analysis. The database ingest, a process called Extract-Transform-Load (ETL), checks that the measurements requested in the ER are fulfilled by each experimental run, and then stores the data and metadata in a database called the data catalog. The Data Converge (DC) tool converts the data catalog records for each experiment into several standardized and versioned data products. These versioned products, such as data frames of log transformed flow cytometry events or per sample absorbance measurements, provide a standard schema across experiments that use different builds, reagents, and protocols. The standardization provides a common format, which enables generic analysis tools that can apply to many different experiments without extensive per-experiment customization. The versioning supports more effective analyst-to-analyst comparisons that are based upon the same version of the data. The RT uses a final tool called the Precomputed Data Table (PDT) (42) to apply a suite of analysis tools to the standardized dataframes. Each analysis tool (described below) processes the data to pre-compute (rather than have the user manually compute) the resulting data. The resulting analyses, in conjunction with any *ad hoc* analyses performed by the user, can be used to inform additional experiments with the RT or inform new designs with the DA components.

In this work, the experimental planning tools automatically launched experimental runs at the Strateos (47) robotic cloud laboratory. Each run produced data that Strateos uploaded to the Texas Advanced Computing Center (TACC) (48) infrastructure for RT processing and analysis. TACC provided the Synergistic Discovery and Design Environment (SD2E; (49)) on which the various components of the RT are integrated. RT packages (and other SD2 packages) are also freely available at (50).

### 4.4. Experimental process

#### 4.4.1. Plasmid construction

Backbone and insert fragments were amplified with PCR, gel extracted, purified, and assembled using Gibson assembly (51) using standardized assembly linkers. Backbones contained a high-copy E. coli origin of replication and ampicillin resistance for propagation. The yeast expression cassettes were flanked upstream and downstream by approximately 500 bases of chromosomal homology to the yeast genome and PmeI restriction sites for linearization before transformations. Plasmids were sequence-verified using Sanger sequencing.

#### 4.4.2. Build construction

Yeast builds were constructed using genomic integration from linearized DNA into the CEN.PK113-7D strain of *S. cerevisae*. Builds were constructed using genomic integration from linearized DNA. Integrative plasmids were linearized using PmeI digestion (37C, 30 min) to cut upstream and downstream of the chromosomal homology. Unpurified, linearized DNA was transformed into yeast cells using a standard lithium acetate protocol (52). Build selection was performed on solid synthetic-complete (SC) using auxotrophic or antibiotic markers. Diagnostic colony PCR was performed to verify integration into the proper locus. Builds were picked from single colonies and stored long-term at -80ºC in a sterilized 30% glycerol and media solution. Build retrieval was performed by plating glycerol stocks onto solid media plates (YPAD) grown for 2-3 days at 30ºC and picking single colonies for liquid culture. All yeast cultures and assays were grown at 30ºC shaking at 275 RPM.

Builds consisted of both individual inducible parts and full circuits. Individual parts were characterized via dosage response flow cytometry experiments to confirm functional response to inducer presence.

#### 4.4.3. Plate layout for circuits

Each 96-well plate had eight data rows, one for each build. The columns had five different inducer concentrations, two replicates for each, as well as two columns for controls. Each plate had one of three media: standard (SC), rich (YEP), or slow (SB) and one logical input transition (see Tables 1-2).

#### 4.4.4. Automated Laboratory

The Strateos robotic cloud lab provides three main functions to DART: (i) the development of replicable, robotically-executed experimental protocols, (ii) the execution of requested experimental runs, and (iii) the delivery of the measurement data to the SD2E platform on TACC.

The protocols developed as part of this and related projects are variations of high-throughput screening with 96-well plates. Each protocol involves commanding a robotic workcell to incubate samples overnight, and then dilute, apply reagents, and gather a time series of plate reader and flow cytometery measurements. Strateos executes experimental runs that it receives from the RT. Each run references a container that includes the samples that include the various designs developed by DA components. Users ship these containers to Strateos and create container metadata describing each sample. The RT’s XPlan planner allocates the requested samples in each ER to the container aliquots using this metadata. The Strateos platform validates each requested experimental run, constructs a sequence of Autoprotocol (53) instructions, and informs the RT of the run identifier. After completing the protocol, the Strateos platform uploads the experimental data and metadata referenced by the run identifier. The RT then ingests the run data as described above.

### 4.5. Analysis tools

#### 4.5.1. Precomputed Data Table (PDT)

The PDT is a component of RT and is responsible for executing a suite of analyses on data products generated by Data Converge (briefly described in Methods 4.3) and storing and versioning the respective results. The PDT provides DART with (i) rapid, consistent analysis of data of different types and from different experiments and (ii) the ability to automatically add additional data features (results from analyses) for higher-level analyses and modeling. These two features enable an increase in the rate at which DART can be iterated as actionable results are available within hours of data availability and more advanced analyses can be applied sooner. The PDT contains several types of analyses and not all are executed on all available data due to incompatibility of analysis types and data types. The next three subsections are descriptions of the analyses that the PDT was programmed to execute for the experiments at hand. The full suite of analyses are listed in (25).

#### 4.5.2. Binary Event Prediction (BEP)

The Binary Event Prediction tool (42; 54) develops a classifier to predict whether an event within a flow cytometry sample corresponds to a “high” or a “low” signal, in the same sense as digital logic. Properly interpreting flow cytometry data collected from cells expressing a fluorescent protein requires the use of a positive and a negative control. In the case of the current experiments, the positive controls are cells that constitutively express GFP and the negative controls are cells that do not express GFP due to the lack of the GFP coding sequence. Using these controls, BEP develops the classifier for each pair of high/positive and low/negative controls. The tool trains the classifier on all flow cytometry events for the high/positive and low/negative controls, labeling each respective event as either 0 (low) or 1 (high), and then uses it to predict for each event within each sample whether the event is low or high.

Compared to techniques that define a linear threshold to a single flow cytometry channel measuring GFP to separate low and high, BEP constructs a multi-dimensional nonlinear threshold (represented as a random forest). The classifier not only subsumes the threshold method, but also incorporates all flow cytometry channels, such as forward scatter, side scatter, and other color channels.

In the redesign effort described above, bimodality was observed in GFP-positive cytometer channels for the positive and negative controls. This caused ambiguity in determining what events are high and low by the classifier as subpopulations of events in one control were consistent with subpopulations in the opposite control. BEP was used to overcome this bimodality in the positive and negative controls. The poor control data were culled using the classifier’s training/test split of the controls during cross-validation to identify the probability of each label predicted in the test set of each cross-validation fold. The control data was then “cleaned” where for each control type, a threshold on probability was applied that dropped events found below the threshold but maintained a minimal total event count of 10,000 events. A new classifier was constructed on the remaining “clean” data and used to predict signals for the non-control samples. The resulting model self-assesses the probability that each control data point should be used in a respective group of labeled points and keeps only the most probable.

#### 4.5.3. Performance Metrics (PM)

Performance Metrics (55) measures the performance of experiments where the samples should be in two different states (e.g. ON and OFF states) and should have a separation in experimental measurements between these two states. The user defines how to aggregate samples (e.g. by build, condition, time) and the experimental measurement (e.g. fluorescence), and the tool computes a suite of metrics that compare the measured values for the two states. The tool returns metrics of how large the separation between the two defined states are for each group in the aggregation.

The metrics compare the distributions of aggregates of samples in each state (ON versus OFF) to compute the differences (ON-OFF) and ratios (ON/OFF) between the two states. The tool compares the left versus right hand side of the distributions via percentiles and standard deviations at different thresholds. For example, PM will aggregate samples by build, condition, and time point, and then for each group, it computes the difference between the 50% percentile of the ON samples and the 50% percentile of the OFF samples. It then computes per sample metrics by comparing each individual sample in one state to the opposite state’s distribution. For example, for each group, for each individual ON sample, it computes the difference and ratio to the median of the OFF states; and then vice versa for each of the OFF samples.

PM reads in a configuration file and data and metadata files. The configuration file specifies the column name of the experiment output, information about the states (e.g. ON and OFF, or any other identifier), as well as a dictionary of what columns to use to make aggregates (e.g. group by build; group by build, condition, time). The per sample metrics for aggregates of build, condition, and time can be used as inputs by Data Diagnosis.

#### 4.5.4. Data Diagnosis (DD)

Data Diagnosis (56) performs diagnostic tests for both experimental design and experimental performance. These test for variables that are associated with variations in performance, and identify which values of the variables are associated with good or bad performance. Additionally, our tests identify if there appears to be any dependence between variables, for example, two variables contain the same information and should not be debugged separately.

DD has two tests for performance and one test for dependence. Our first performance test is the Kruskal-Wallis H-test for categorical variables. This completes a non-parametric analysis of variance for the categorical variables grouped by a user selected variable. The continuous variable analysis uses the Spearman correlation coefficient to measure the association between the performance variable and each continuous variable. Finally, there is the dependence test for randomization. This is primarily to study whether the experimental design has redundancies or has missed combinations of variables. It should be run prior to the experiment, but can also be run afterwards to aid in analysis and troubleshooting of the experiment. This test determines if the dependent variables were properly randomized by running a Chi-Squared Test for Independence between all pairs of categorical features. If the experiment was properly randomized then none of the pairs should be dependent. Together all of these tests provide researchers valuable information on how their experiment performed, and most importantly if certain variables are associated with good or poor performance.

DD reads in a configuration file and data and metadata files. The configuration file specifies the column name of the performance values (in this case, coming from performance metrics), as well as a dictionary of what columns to use to perform analysis on subsets of the metadata.

#### 4.5.5. Parameter Value Estimations and ODE Modeling

A data-fitting algorithm using a Nelder–Mead minimization method (57) was implemented to determine the Hill function (58) parameter values for the induction and repression dynamics of the different genetic parts used in the OR/NOR circuit designs. The experimental data used for the fitting algorithm was obtained from the geometric mean of flow cytometry data distributions from the dosage response experiments as described in section 4.4.2. The experimental data were fitted to Equation 1 (for activations), and Equation 2 (for repressions) derived from Cello (16):

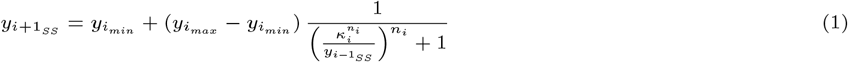

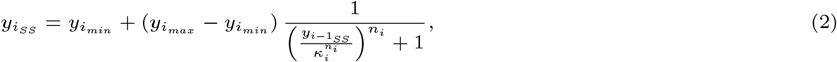

where *y*_*iss*_ is the steady-state output promoter activity of part *i; y_imin_* and *y*_*imax*_ are the minimal and maximal output promoter activities (obtained from experimental results), respectively, for part *i*; *κ*_*i*_ and *n*_*i*_ are the affinity and cooperativity of transcription factor binding (obtained with the fitting algorithm); and, finally, *y*_*i−*1*SS*_ is the steady-state input promoter activity from the previous part’s output (calculated also using Equations 1 or 2). Using the Hill function parameter value estimations a resulting ODE model is then analyzed using the Runge-Kutta-Fehlberg (4,5) method (59) implemented in iBioSim (60) to obtain steady-state output predictions for each design under different input concentrations (shown in Appendix Figs. 14 and 15).

## 5. Data availability

Processed data and scripts used in the generation of figures in this manuscript are available in a public repository. The code packages cited in the manuscript are all open source.

## 6. Competing interests

Some of the authors are employed by companies that may benefit or be perceived to benefit from this publication.

## 7. Author contributions statement

Conceptualization: BC, JV, RCM, MG, TG, KM, SBH.

Data curation: BC, JV, RCM, AD, DB, TJ, MW, GZ, JN, CJM, JB, BB, TM, TTN, NR. Formal analysis: BC, RCM, AD, PF, TJ, MW, GZ, KLJ, RPG.

Funding acquisition: DB, MWV, CJM, TG, KM, SBH. Investigation: JV, RCM, JN, VB, TRH, LAM.

Methodology: BC, JV, RCM, HE, AD, PF, FCM, DB, ME, KLJ, MG, JB.

Project administration: DB, JB, MW, MWV, SBH. Resources: JV, JN, NG, MWV, JU.

Software: BC, JV, RCM, HE, AD, PF, FCM, DB, TJ, MW, GZ, ME, SG, RPG, CJM, MG, NG, JU, JB, BB, TM, TTN, NR.

Supervision: DB, JB, MW, TG, KM, SBH. Validation: JV, AD, MW, GZ, JN.

Visualization: BC, RCM, HE, AD, PF, MW, KLJ.

Writing, original draft: BC, JV, RCM, AD, PF, DB, ME, KLJ, CJM.

Writing, review and editing: BC, PF, DB, FCM, ME, RPG, JB, TG, KM, SBH.

## 8. Acknowledgments

This work was supported by Air Force Research Laboratory (AFRL) and DARPA contracts HR001117C0092, HR001117C0094, HR001117C0095, FA875017C0184, FA875017C0229, FA875017C0231, and FA875017C0054 as part of the Synergistic Discovery and Design (SD2) program. This document does not contain technology or technical data controlled under either U.S. International Traffic in Arms Regulation or U.S. Export Administration Regulations. Views, opinions, and/or findings expressed are those of the author(s) and should not be interpreted as representing the official views or policies of the Department of Defense or the U.S. Government.

## A. Experiment-to-design keys

**Fig. 13.**
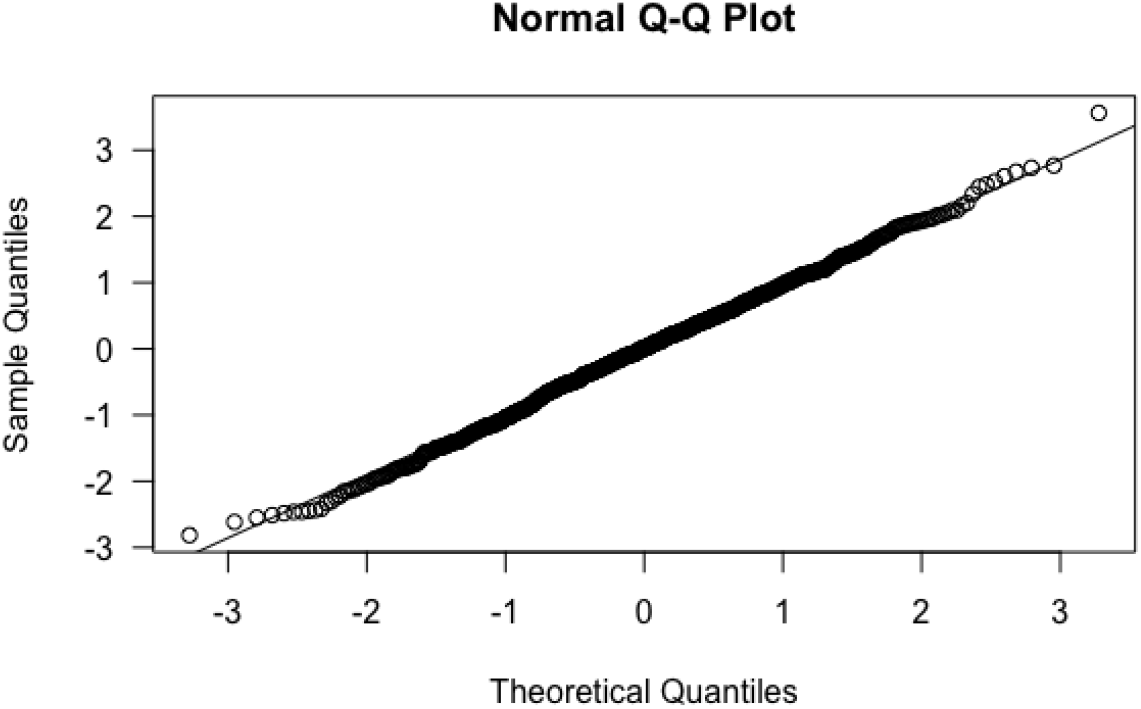
Normal Quantile-Quantile plot with quantile residuals for logistic regression model

## B. Binary Event Prediction

## C. Fold change of the BEP cleaned data compared with unmodified data

For technical reasons in the software, the plates were split between those that titrated a single inducer (Titration, 12 plates) and those that required simultaneous addition of both inducers (Fixed Ratio, 3 plates). The figures in this section are split by these two experiments. Fig. 10 is the output of Performance Metrics and Fig. 11-12 are produced by Data Diagnosis. The BEP cleaned data are the subset of the log-transformed unmodified data associated to the greater BEP ratio.

As seen in Fig. 10, fold changes hover around 1, likely due to strong bimodality. Median fold changes less than 1 indicate circuit failures. The BEP clean model does not increase median fold change. The primary difference induced by BEP cleaning is the production of outliers with high fold change that skew the average.

Fig. 11 and 12 show the variance in fold change as a function of different variables, including the OR or NOR circuit (`build_circuit`), the design (`build`, see Table 3), plate (exp`eriment_id`, see Table 2), media condition, and time point. The remaining categories have difficulties with their interpretation due to the aggregations involved. Fig. 11 shows the analysis for the log-transformed unmodified data and Fig. 12 shows the analysis for the BEP cleaned data. The unmodified data show slight differences in performance in circuit and design, but for the BEP-cleaned data these patterns are swamped by large outliers in fold change.

**Table 3.** Key relating experimental build identifiers to the design nomenclature as seen in Fig. 11 and 12.

| Build Identifier | Design |
| --- | --- |
| UWBF_36564 | DSGRN NOR, CDM low |
| UWBF_36565 | DSGRN NOR, CDM high |
| UWBF_36566 | DSGRN OR, CDM low |
| UWBF_36567 | DSGRN OR, CDM high |
| UWBF_36568 | Simple NOR, CDM low |
| UWBF_36569 | Simple NOR, CDM high |
| UWBF_36570 | Simple OR, CDM low |
| UWBF_36571 | Simple OR, CDM high |

## D. Logistic regression model and diagnostics

In order to assess the impact of each variable on the probability of success and produce estimated probabilities of success, a logistic regression model was fit. To account for the dependencies between plates, first a mixed effects model was fit with a random intercept for the plate variable. Since the variance of the random intercepts was negligibly different from zero, the model was refit without the random effect for plate. As the removal of the random effect allows for models accounting for *overdispersion*, variability in the responses beyond what is expected under the binomial distribution, a second model was fit with the overdispersion parameter estimated as 1.016. Since the overdispersion parameter is one in the absence of overdispersion, a final model was fit without allowing for overdispersion.

This model included interaction effects for circuit with both topology and CDM low/high design in order to test whether the effect of topology or CDM low/high design depends on the circuit. The estimated model is as follows:

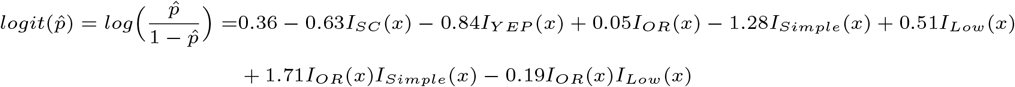

where *I*_*SC*_ (*X*) is an indicator function with *I*_*SC*_ (*X*) = 1 if the observation, *x*, is in the SC media category and *I*_*SC*_ (*X*) = 0 otherwise. Note that the estimates can be exponentiated to produce estimates of the multiplicative effects on the odds of success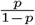 The table of estimates with p-values and 95% confidence intervals are below.

There is very strong evidence that the effect of circuit depends on topology (p-value*<*0.0001) and very weak evidence that the effect of circuit depends on CDM prediction (p-value=0.4865). A Normal Q-Q plot using quantile residuals, which should be normally distributed with the choice of an appropriate link function, is included below and indicated no issues with the logit link function. This link function transforms the probability of success, which does not have a linear relationship with the predictors, to one that does: the log odds of success.

**Table 4.**
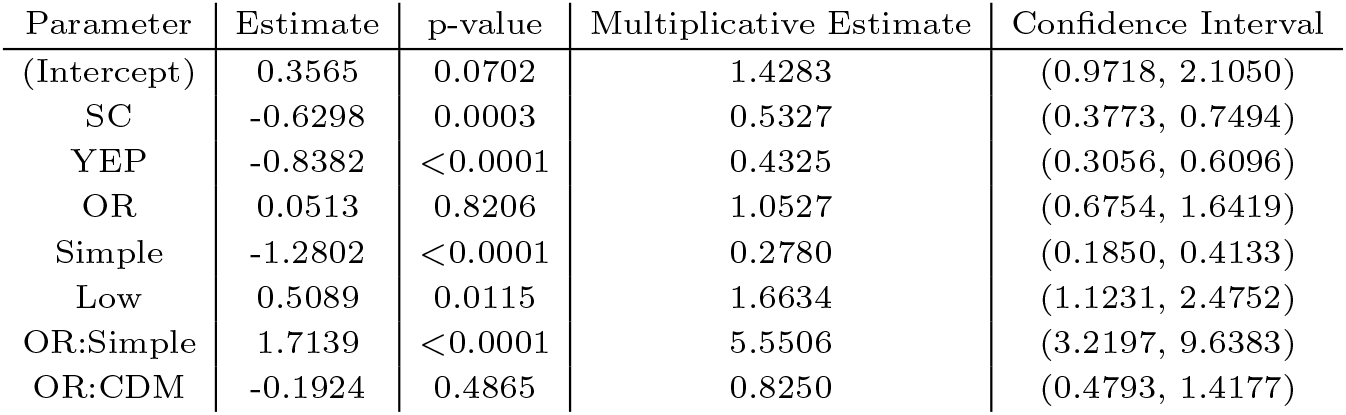
Estimates, p-values, and confidence intervals for logistic regression model.

## E. Comprehensive results for ODE simulations

**Fig. 14.**
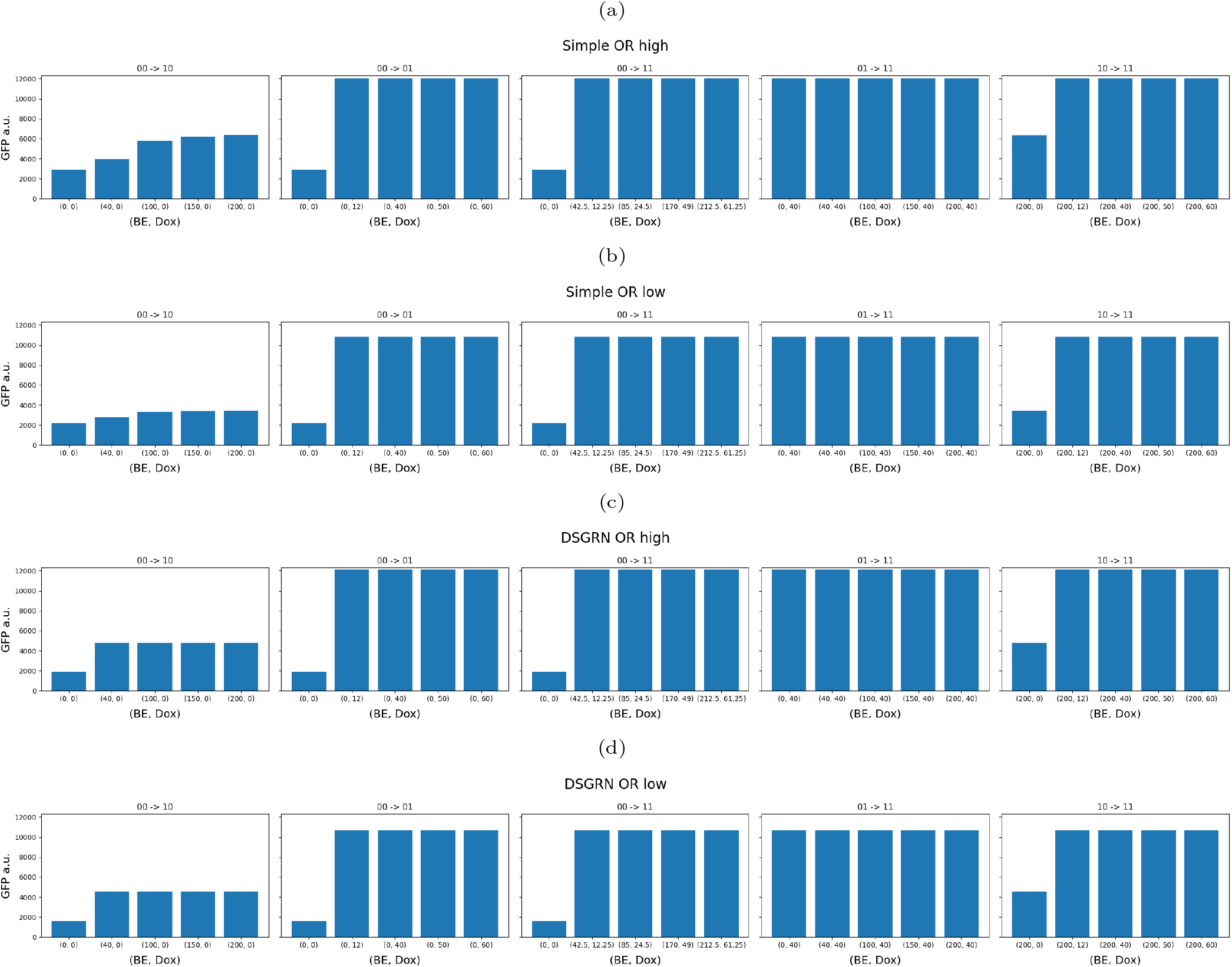
Predicted steady-state values of the geometric mean of the flow cytometry distribution of GFP a.u. using estimated Hill function parameter values for: (a) Simple OR/CDM high design, (b) Simple OR/CDM low design, (c) DSGRN OR/CDM high design, and (d) DSGRN OR/CDM low design. An OFF circuit state corresponds to approximately 1500-2000 a.u.; robust ON circuit states occur at about 8000 a.u. and up, and weak ON states occur at about 4000 a.u. Columns 1-3 (transitions 00 → 10, 00 → 01, 00 → 11): The left-most bar in each bar graph should be OFF. All other bars should be ON. Columns 4-5 (transitions 01 → 11, 10 → 11): All bars should be ON.

**Fig. 15.**
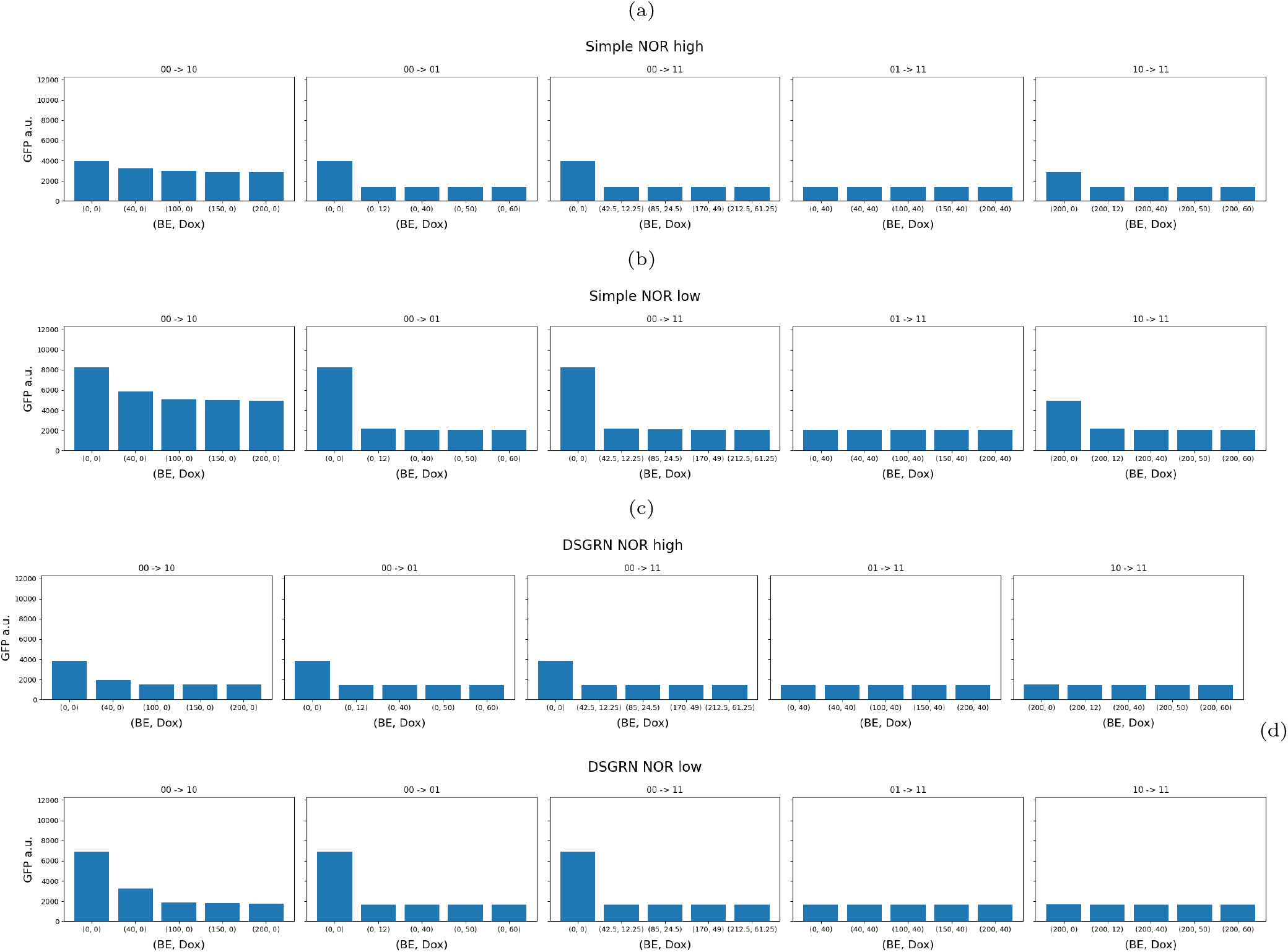
Predicted steady-state values of the geometric mean of the flow cytometry distribution of GFP a.u. using estimated Hill function parameter values for: (a) Simple NOR/CDM low design, (b) Simple NOR/CDM low design, (c) DSGRN NOR/CDM high design, and (d) DSGRN NOR/CDM low design. An OFF circuit state corresponds to approximately 1500-2000 a.u.; robust ON circuit states occur at about 8000 a.u. and up, and weak ON states occur at about 4000 a.u. Columns 1-3 (transitions 00 → 10, 00 → 01, 00 → 11): The left-most bar in each bar graph should be ON. All other bars should be OFF. Columns 4-5 (transitions 01 → 11, 10 → 11): All bars should be OFF.

## References

1. I. Del Valle, E. M. Fulk, P. Kalvapalle, J. J. Silberg, C. A. Masiello, and L. B. Stadler, “Translating new synthetic biology advances for biosensing into the earth and environmental sciences,” Frontiers in Microbiology, p. 3513, 2021.

2. A. S. Khalil and J. J. Collins, “Synthetic biology: applications come of age,” Nature Reviews Genetics, vol. 11, no. 5, pp. 367–379, 2010.

3. D. E. Cameron, C. J. Bashor, and J. J. Collins, “A brief history of synthetic biology,” Nature Reviews Microbiology, vol. 12, no. 5, pp. 381–390, 2014.

4. B. Cummins, R. C. Moseley, A. Deckard, M. Weston, G. Zheng, D. Bryce, S. Gopaulakrishnan, T. Johnson, J. Nowak, M. Gameiro, T. Gedeon, K. Mischaikow, M. Vaughn, N. I. Gaffney, J. Urrutia, R. P. Goldman, J. Beal, B. Bartley, T. T. Nguyen, N. Roehner, T. Mitchell, J. D. Vrana, K. J. Clowers, N. Maheshri, D. Becker, E. Mikhalev, V. Biggers, T. R. Higa, L. A. Mosqueda, and S. B. Haase, “Computational prediction of synthetic circuit function across growth conditions.”

5. M. A. Marchisio and J. Stelling, “Computational design tools for synthetic biology,” Current opinion in biotechnology, vol. 20, no. 4, pp. 479–485, 2009.

6. A. Villalobos, J. E. Ness, C. Gustafsson, J. Minshull, and S. Govindarajan, “Gene designer: a synthetic biology tool for constructing artificial DNA segments,” BMC bioinformatics, vol. 7, no. 1, pp. 1–8, 2006.

7. N. Swainston, M. Dunstan, A. J. Jervis, C. J. Robinson, P. Carbonell, A. R. Williams, J.-L. Faulon, N. S. Scrutton, and D. B. Kell, “Partsgenie: an integrated tool for optimizing and sharing synthetic biology parts,” Bioinformatics, vol. 34, no. 13, pp. 2327–2329, 2018.

8. M. J. Czar, Y. Cai, and J. Peccoud, “Writing DNA with genoCAD™,” Nucleic acids research, vol. 37, no. uppl 2, pp. W40–W47, 2009.

9. G. Misirli, J. S. Hallinan, T. Yu, J. R. Lawson, S. M. Wimalaratne, M. T. Cooling, and A. Wipat, “Model annotation for synthetic biology: automating model to nucleotide sequence conversion,” Bioinformatics, vol. 27, no. 7, pp. 973–979, 2011.

10. L. Huynh, A. Tsoukalas, M. Köppe, and I. Tagkopoulos, “Sbrome: A scalable optimization and module matching framework for automated biosystems design,” ACS Synthetic Biology, vol. 2, no. 5, pp. 263–273, 2013, pMID: 23654271. [Online]. Available: https://doi.org/10.1021/sb300095m

11. Y. Park, A. Espah Borujeni, T. E. Gorochowski, J. Shin, and C. A. Voigt, “Precision design of stable genetic circuits carried in highly-insulated e. coli genomic landing pads,” Molecular systems biology, vol. 16, no. 8, p. e9584, 2020.

12. P. François and V. Hakim, “Design of genetic networks with specified functions by evolution in silico,” Proceedings of the National Academy of Sciences, vol. 101, no. 2, pp. 580–585, 2004.

13. U. Kuter et al., “XPLAN: Experiment planning for synthetic biology,” in ICAPS Workshop on Hierarchical Planning, 2018.

14. J. Vrana, O. de Lange, Y. Yang, G. Newman, A. Saleem, A. Miller, C. Cordray, S. Halabiya, M. Parks, E. Lopez et al., “Aquarium: open-source laboratory software for design, execution and data management,” Synthetic Biology, vol. 6, no. 1, p. ysab006, 2021.

15. T. S. Jones, S. Oliveira, C. J. Myers, C. A. Voigt, and D. Densmore, “Genetic circuit design automation with cello 2.0,” Nature Protocols, pp. 1–17, 2022.

16. A. A. K. Nielsen, B. S. Der, J. Shin, P. Vaidyanathan, V. Paralanov, E. A. Strychalski, D. Ross, D. Densmore, and C. A. Voigt, “Genetic circuit design automation,” Science, vol. 352, no. 6281, p. aac7341, Apr. 2016.

17. T. Schladt, N. Engelmann, E. Kubaczka, C. Hochberger, and H. Koeppl, “Automated design of robust genetic circuits: Structural variants and parameter uncertainty,” ACS synthetic biology, vol. 10, no. 12, pp. 3316–3329, 2021.

18. E. Appleton, C. Madsen, N. Roehner, and D. Densmore, “Design automation in synthetic biology,” Cold Spring Harbor perspectives in biology, vol. 9, no. 4, p. a023978, 2017.

19. J. W. Yeoh, N. Swainston, P. Vegh, V. Zulkower, P. Carbonell, M. B. Holowko, G. Peddinti, and C. L. Poh, “SynBiopython: an open-source software library for Synthetic Biology,” Synthetic Biology, vol. 6, no. 1, 02 2021, ysab001. [Online]. Available: https://doi.org/10.1093/synbio/ysab001

20. B. Cummins, T. Gedeon, S. Harker, K. Mischaikow, and K. Mok, “Combinatorial representation of parameter space for switching networks,” SIAM Journal on Applied Dynamical Systems, vol. 15, no. 4, pp. 2176–2212, 2016. [Online]. Available: http://epubs.siam.org/doi/abs/10.1137/15M1052743

21. E. Yeung, J. Kim, Y. Yuan, J. Gonçalves, and R. M. Murray, “Data-driven network models for genetic circuits from time-series data with incomplete measurements,” Journal of the Royal Society Interface, vol. 18, no. 182, p. 20210413, 2021.

22. E. Yeung, S. Kundu, and N. Hodas, “Learning deep neural network representations for koopman operators of nonlinear dynamical systems,” in 2019 American Control Conference (ACC). IEEE, 2019, pp. 4832–4839.

23. Z. Li and Q. Yang, “Systems and synthetic biology approaches in understanding biological oscillators,” Quantitative Biology, vol. 6, no. 1, pp. 1–14, 2018.

24. M. M. Jessop-Fabre and N. Sonnenschein, “Improving reproducibility in synthetic biology,” Frontiers in Bioeng. and Biotech., vol. 7, pp. 1–18, 2019. [Online]. Available: https://www.frontiersin.org/article/10.3389/fbioe.2019.00018

25. D. Bryce, R. P. Goldman,, M. DeHaven, J. Beal, B. Bartley, T. T. Nguyen, N. Walczak, M. Weston, G. Zheng, J. Nowak, P. Lee, J. Stubbs, N. Gaffney, M. W. Vaughn, C. J. Myers, R. C. Moseley, S. Haase, A. Deckard, B. Cummins, and N. Leiby, “Round Trip: An Automated Pipeline for Experimental Design, Execution, and Analysis,” ACS Syn. Bio., 2022. [Online]. Available: https://doi.org/10.1021/acssynbio.1c00305

26. D. Bryce, R. P. Goldman, M. Dehaven, J. Beal, T. Nguyen, N. Walczak, M. Weston, G. Zheng, J. Nowak, J. Stubbs, M. Vaughn, N. Gaffney, and C. Myers, “Round-trip: An automated pipeline for experimental design, execution, and analysis,” in Proceedings of the 12th International Workshop on Bio-Design Automation (IWBDA-20), 2020, pp. 29–30.

27. N. Roehner et al., “Sharing structure and function in biological design with sbol 2.0,” ACS Syn. Bio., vol. 5, no. 6, pp. 498–506, 2016. [Online]. Available: https://doi.org/10.1021/acssynbio.5b00215

28. N. Roehner, J. Mante, C. J. Myers, and J. Beal, “Synthetic biology curation tools (synbict),” ACS Synthetic Biology, vol. 10, no. 11, pp. 3200–3204, 2021.

29. M. Gameiro, “DSGRN software,” 2022. [Online]. Available: https://github.com/marciogameiro/DSGRN

30. M. Gameiro, T. Gedeon, S. Kepley, and K. Mischaikow, “Rational design of complex phenotype via network models,” PLoS computational biology, vol. 17, no. 7, p. e1009189, 2021.

31. M. Eslami, A. E. Borujeni, H. Doosthosseini, M. Vaughn, H. Eramian, K. Clowers, D. B. Gordon, N. Gaffney, M. Weston, D. Becker, Y. Dorfan, J. Fonner, J. Urrutia, C. Corbet, G. Zheng, J. Stubbs, A. Cristofaro, P. Maschhoff, J. Singer, C. A. Voigt, and E. Yeung, “Prediction of whole-cell transcriptional response with machine learning,” bioRxiv, 2021. [Online]. Available: https://www.biorxiv.org/content/early/2021/05/01/2021.04.30.442142

32. H. Eramian and M. Eslami, “Combinatorial design model,” 2021. [Online]. Available: https://github.com/SD2E/CDM

33. J. Vrana, “DASi DNA design,” 2021. [Online]. Available: https://github.com/jvrana/DASi-DNA-Design.git

34. J. Vrana, “DASi DNA design documentation,” 2021. [Online]. Available: https://jvrana.github.io/DASi-DNA-Design/

35. J. Vrana, “Software systems for automated manufacturing of engineered organisms,” Ph.D. dissertation, University of Washington, 2021, copyright - Database copyright ProQuest LLC; ProQuest does not claim copyright in the individual underlying works; Last updated - 2021-11-24. [Online]. Available: https://www.proquest.com/dissertations-theses/software-systems-automated-manufacturing/docview/2594492440/se-2?accountid=28148

36. J. Vrana, “Terrarium,” 2021. [Online]. Available: https://github.com/jvrana/Terrarium.git

37. “Aquarium: The laboratory operating system,” https://www.aquarium.bio/, accessed: 2022-05-09.

38. B. Cummins, “DSGRN design interface software,” 2021. [Online]. Available: https://gitlab.com/breecummins/dsgrndesigninterface.git

39. R. P. Goldman, R. Moseley, N. Roehner, B. Cummins, J. D. Vrana, K. J. Clowers, D. Bryce, J. Beal, M. DeHaven, J. Nowak, T. Higa, V. Biggers, P. Lee, J. P. Hunt, S. B. Haase, M. Weston, G. Zheng, A. Deckard, S. Gopaulakrishnan, J. F. Stubbs, N. I. Gaffney, M. W. Vaughn, N. Maheshri, E. Mikhalev, B. Bartley, R. Markeloff, T. Mitchell, T. Nguyen, D. Sumorok, N. Walczak, C. Myers, Z. Zundel, B. Hatch, J. Scholz, and J. Colonna-Romano, “Highly-automated, high-throughput replication of yeast-based logic circuit design assessments,” bioRxiv, 2022. [Online]. Available: https://www.biorxiv.org/content/early/2022/06/01/2022.05.31.493627

40. M. W. Gander, J. D. Vrana, W. E. Voje, J. M. Carothers, and E. Klavins, “Digital logic circuits in yeast with CRISPR-dCas9 NOR gates,” Nature Communications, vol. 8, 2017. [Online]. Available: http://www.ncbi.nlm.nih.gov/pmc/articles/PMC5458518/

41. S. Kepley, K. Mischaikow, and E. Queirolo, “Global analysis of regulatory network dynamics: equilibria and saddle-node bifurcations,” 2022. [Online]. Available: https://arxiv.org/abs/2204.13739

42. G. Zheng, R. C. Moseley, D. Bryce, A. Deckard, B. Cummins, R. Goldman, H. Eramian, T. Johnson, and M. Weston, “Pre-computed data table,” 2021. [Online]. Available: https://github.com/SD2E/precomputed-data-table.git

43. S. Lundberg, “Shapley additive explanations (shap),” 2018. [Online]. Available: https://github.com/slundberg/shap

44. T. Gedeon, B. Cummins, S. Harker, and K. Mischaikow, “Identifying robust hysteresis in networks,” PLOS Computational Biology, vol. 14, no. 4, p. e1006121, 2018. [Online]. Available: https://journals.plos.org/ploscompbiol/article?id=10.1371/journal.pcbi.1006121

45. “University of Washington Biofabrication Center,” http://www.uwbiofab.org/, accessed: 2022-05-06.

46. T. Nguyen, N. Walczak, D. Sumorok, M. Weston, and J. Beal, “Intent parser: A tool for codification and sharing of experimental design,” ACS Synthetic Biology, vol. 11, no. 1, pp. 502–507, 2021.

47. “Strateos, cloud lab automation-as-a-service,” https://strateos.com, accessed: 2022-05-06.

48. “Texas Advanced Computing Center,” https://www.tacc.utexas.edu/, accessed: 2022-05-06.

49. “Synergistic discovery and design environment,” https://www.tacc.utexas.edu/research-development/tacc-projects/sd2e, accessed: 2022-05-23.

50. “Synergistic discovery and design github,” https://github.com/SD2E, accessed: 2022-05-23.

51. D. G. Gibson, L. Young, R.-Y. Chuang, J. C. Venter, C. A. Hutchison, and H. O. Smith, “Enzymatic assembly of dna molecules up to several hundred kilobases,” Nature methods, vol. 6, no. 5, pp. 343–345, 2009.

52. R. D. Gietz and R. A. Woods, “Transformation of yeast by lithium acetate/single-stranded carrier dna/polyethylene glycol method,” in Methods in enzymology. Elsevier, 2002, vol. 350, pp. 87–96.

53. “Autoprotocol: An open standard for scientific experimental design and automation.” https://autoprotocol.org, accessed: 2022-05-10.

54. D. Bryce, R. Moseley, and J. Ladwig, “Python sd2 circuit analysis tool,” 2022. [Online]. Available: https://github.com/SD2E/pysd2cat.git

55. A. Deckard and T. Johnson, “Performance metrics,” 2019. [Online]. Available: https://github.com/SD2E/performance-metrics.git

56. A. Deckard and T. Johnson, “Data diagnosis,” 2019. [Online]. Available: https://github.com/SD2E/diagnose.git

57. J. A. Nelder and R. Mead, “A simplex method for function minimization,” The computer journal, vol. 7, no. 4, pp. 308–313, 1965.

58. M. Santillàn, “On the use of the hill functions in mathematical models of gene regulatory networks,” Mathematical Modelling of Natural Phenomena, vol. 3, no. 2, pp. 85–97, 2008.

59. E. Fehlberg, “Low-order classical Runge-Kutta formulas with stepsize control and their application to some heat transfer problems,” National aeronautics and space administration, Tech. Rep., Jul. 1969, 00572. [Online]. Available: https://ntrs.nasa.gov/search.jsp?R=19690021375

60. L. Watanabe, T. Nguyen, M. Zhang, Z. Zundel, Z. Zhang, C. Madsen, N. Roehner, and C. Myers, “iBioSim 3: A tool for model-based genetic circuit design,” ACS Synthetic Biology, Jun. 2018. [Online]. Available: https://doi.org/10.1021/acssynbio.8b00078

